# Diversifying selection and adaptive introgression of carotenoid-processing genes underlie the evolution of bill color in the long-tailed finch

**DOI:** 10.1101/2024.06.17.599356

**Authors:** Daniel M. Hooper, Callum S. McDiarmid, Matthew J. Powers, Nicholas M. Justyn, Marek Kučka, Nathan S. Hart, Geoffrey E. Hill, Peter Andolfatto, Yingguang Frank Chan, Simon C. Griffith

## Abstract

Carotenoid pigmentation produces the yellow and red coloration of birds and other vertebrates, but our understanding of the genetic architecture of carotenoid ornamentation is largely limited to studies of novel color variants observed in captively bred populations. The complexity of carotenoid-based color evolution in nature remains poorly characterized. Here, we examine the long-tailed finch *Poephila acuticauda*, an Australian songbird with two hybridizing subspecies that differ in bill coloration: yellow in western subspecies *acuticauda* and red in eastern subspecies *hecki*. We characterize the carotenoid composition of each subspecies and find that yellow bills can be explained by the loss of C(4)-oxidation, thus blocking yellow dietary pigments from being metabolized to red. Combining linked-read genomic sequencing and reflectance spectrophotometry measurements of bill color collected from wild-sampled finches and laboratory crosses, we identify four loci that together explain 53% of variance in this trait. The two loci of largest effect contain the genes *CYP2J19*, an essential enzyme for the ketolation via C(4)-oxidation of dietary carotenoids, and *TTC39B*, an enhancer of ketocarotenoid production. Evolutionary genealogy reconstruction indicates that the red-billed phenotype is ancestral and yellow alleles at both *CYP2J19* and *TTC39B* arose and fixed in *acuticauda* approximately 100 kya. Yellow alleles then introgressed into *hecki* less than 5 kya. Across all four loci, *acuticauda* derived variants show evidence of selective sweeps, implying that yellow bill coloration has been favored by natural selection. Our study suggests that the frequent adaptive evolutionary transitions between red and yellow ornamentation in nature can have a simple genetic basis.

**Significance:** We studied variation in carotenoid ornamentation of an Australian songbird with two hybridizing subspecies that differ in bill color: one yellow and the other red. We identified a single metabolic process, C(4)-oxidation, underlying the distinct carotenoid composition of these two bill colors. Genetic association mapping revealed four major effect loci that explained most of the observed variation the trait, including the oxidative ketolation enzyme *CYP2J19* and the carotenoid ketolation enhancer gene *TTC39B*. Evolutionary reconstruction indicates that yellow alleles are derived, ancient (~100 kya), and under positive selection. This has driven their recent (<5 kya) adaptive introgression across the hybrid zone. These findings have important implications for understanding the role of natural selection in phenotypic evolution in natural systems.

## Introduction

Understanding the diversity of color displays in animals stands as a major challenge in evolutionary biology (Cuthill et al. 2017; Price 2017). The yellow, orange, and red colors that regularly adorn birds, fish, and reptiles are among the most conspicuous visual signals of animals. Studies have shown that carotenoid-based coloration plays important and varied roles in social and sexual selection (Svensson and Wong 2011; Hill and McGraw 2006). As such, carotenoid coloration has been proposed to contribute to speciation via the establishment or maintenance of reproductive isolation – for example via species recognition and mating preferences (West-Eberhard 1983; Price 1998; Seddon et al., 2013; Gomes et al., 2016; Price-Waldman et al., 2020).

Despite the importance of animal coloration in evolution, researchers have only recently gained insights into the genetic mechanisms underlying the diversity of carotenoid ornamentation among animals (Toews et al., 2017; Price-Waldman and Stoddard 2021). It has long been known that the carotenoid pigments used to produce red and yellow coloration cannot be synthesized by most vertebrates *de novo* but rather are sourced from their diets (Hill and McGraw 2006; Sefc et al., 2014; Maoka 2020). Crucially, most of the carotenoid pigments ingested by vertebrates are yellow, so red carotenoid coloration requires a metabolic conversion. In the last decade, a combination of experimental and field-based studies has identified a small set of genes essential to produce yellow and red carotenoid coloration. Key among these newly discovered genes is 3-hydroxybutyrate dehydrogenase 1-like (*BDH1L*) and cytochrome P450 2J19 (*CYP2J19*). When expressed without *CYP2J19*, *BDH1L* produces yellow ε,ε-carotenoid pigments such as canary xanthophylls from dietary carotenoids like lutein and zeaxanthin (Toomey et al., 2022a). When *CYP2J19* and *BDH1L* are expressed together, these same dietary carotenoids are metabolized to ketocarotenoids, a group of carotenoids responsible for red coloration (Mundy et al., 2016; Lopes et al., 2016; Toomey et al., 2022a).

While we have a growing understanding of the key enzymes involved in the production of yellow and red coloration in birds and other vertebrates, most of these recent insights have been drawn from studies of domesticated lineages subject to intense artificial selection for novel color variation. For example, scavenger receptor B1 (*SCARB1*), a key carotenoid transport gene, was discovered in a “white recessive” canary breed with entirely white plumage and a congenital vitamin A deficiency due to splice donor site mutation in *SCARB1* (Toomey et al., 2017). This mutation is also associated with a loss of female mating preferences for ornamental coloration (Koch and Hill 2019). The role of tetracopeptide repeat protein 39B (*TTC39B*) as an enhancer of carotenoid ketolation was identified in orange-mutant red-throated parrotfinches *Erythrura psittacea*. This novel morph, created by bird fanciers, develop orange instead of red plumage and plausibly experience impaired color vision due to a splice altering duplication within *TTC39B* which compromises ketocarotenoid production in feathers and retinal cone photoreceptors (Toomey et al., 2022a). The two studies that first identified the essential role of *CYP2J19* in carotenoid ketolation also used color mutants not found in nature (Mundy et al., 2016; Lopes et al., 2016). The *yellowbeak* morph of the zebra finch *Taeniopygia guttata* has yellow bills and tarsi rather than the wildtype red due to a deletion of a functional *CYP2J19* copy that renders them unable to synthesize red ketocarotenoids (Mundy et al. 2016). In “red factor” canaries, bird fanciers purposefully moved a “red factor” (a *CYP2J19* allele) from a red-feathered species, the red siskin *Spinus cucullatus*, to yellow common canaries *Serinus canaria* via hybridization and serial backcrossing (Lopes et al., 2016). In both finches and canaries, artificial selection overcame what would have been negative sexual selection because the novel coloration made these individuals less attractive as mates (Simons and Verhulst 2011; Koch and Hill 2019). Thus, the causative mechanisms underlying carotenoid-based color variation identified from these avicultural forms are often highly pleiotropic mutations and primarily loss-of-function mutations that would be deleterious to carriers in the wild. Almost nothing is currently known about how carotenoid-based traits evolve in nature, the role of species boundaries in the divergence of color traits, and the respective roles of natural and sexual selection in the divergence of carotenoid ornamentation between species (Toews et al., 2017; Price-Waldman and Stoddard 2021).

The long-tailed finch *Poephila acuticauda* is a songbird endemic to the northern tropics of Australia and provides a unique opportunity to study the evolution of carotenoid-based ornamental coloration in a natural system. It consists of two subspecies that differ prominently in bill coloration: yellow in western subspecies *acuticauda* and red in eastern subspecies *hecki* (Fig. 1A and 1B). The two subspecies form a hybrid zone at the edge of the Kimberley Plateau in Western Australia. Notably, the transition from yellow to red ornamentation is displaced relative to the genomic hybrid zone: the center of bill coloration admixture is located ~350 km to the east of this within the Northern Territory (Griffith and Hooper 2017; Hooper et al., 2019). This displacement between the centers of genomic and bill color admixture could be the result of asymmetrical introgression of *acuticauda* color alleles into an otherwise *hecki* genetic background. This makes the long-tailed finch an attractive system to study how natural and sexual selection shape carotenoid ornamentation and the role of color displays as reproductive barriers between species.

**Fig. 1.**
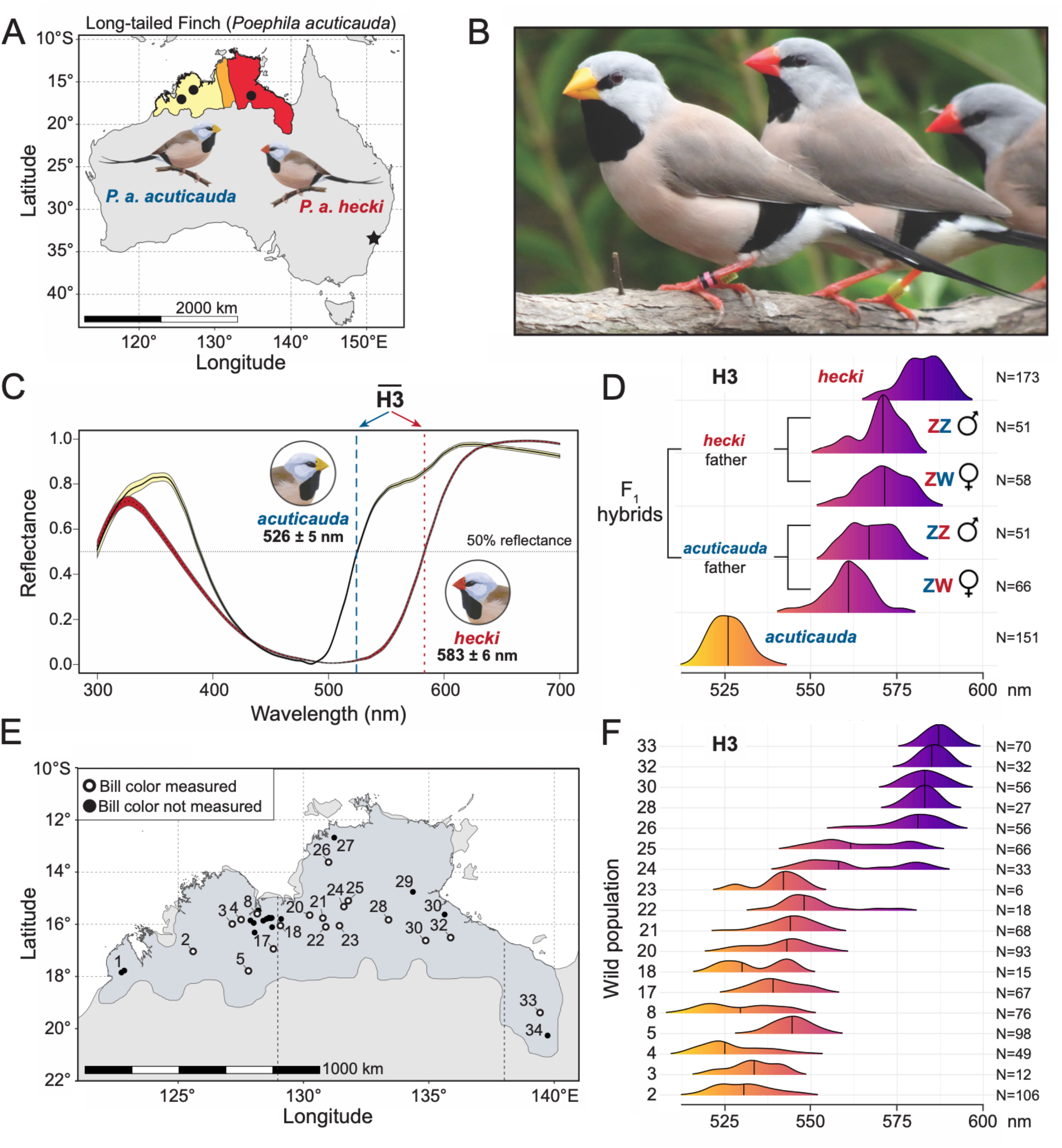
Bill color differentiation between subspecies of the long-tailed finch. (A) Approximate geographic distribution of bill color variation across the range of the long-tailed finch. Western populations have subspecies *acuticauda*-type yellow bills, eastern populations have subspecies *hecki*-type red bills, and a region of phenotypic admixture with individuals bearing orange bills is located in between. The source locations for birds used to establish the captive research colony at Macquarie University in Sydney, NSW, are represented as black circles and a black star, respectively (see Materials and Methods for exact locations). (B) Representative photograph taken of long-tailed finches of each subspecies from our research colony: from left, subspecies *acuticauda*, subspecies *hecki*. (C) Standardized reflectance spectra for bills from individuals of subspecies *acuticauda* (N = 71) and *hecki* (N = 80) reared in common garden conditions and measured via UV-vis reflectance spectrophotometry are shown as mean (solid line) and standard error (yellow and red shading, respectively). The mean and standard deviation for bill hue (colorimetric variable H3 or λ_R50_; the wavelength midway between maximum and minimum reflectance between 400 and 700 nm) is given for each subspecies with vertical dashed lines representing the population mean. (D) Variation in bill hue (H3) observed for parental individuals of both subspecies and their first generation (F_1_) hybrids. Hybrids are grouped by the direction of hybrid cross and sex (ZZ: males; ZW: females). (E) Geographic distribution of populations sampled in this study across the range of the long-tailed finch (shown in blue). Of the 34 populations sampled, the 18 where bill color was measured via reflectance spectrophotometry are shown as hollow circles. (F) Variation in bill hue (H3) observed across the range of the long-tailed finch. Populations arranged from bottom to the top along our west-to-east transect across the range of the species. The number of individuals measured in each population is shown to the right of each ridgeline plot. A total of 948 individuals were measured in the wild.

Our investigation involves four steps. First, we describe the distinct carotenoid composition of the bills of each subspecies and screen for differences in the retinal cone cells of the subspecies using birds reared in captivity under common garden conditions. Second, we examine the genetic architecture of bill color variation using 508 phenotyped individuals from across nearly the entire wild range of the species. Third, we utilize geographic and genomic cline analyses to evaluate evidence of introgression of bill color alleles between subspecies. Fourth, we leverage population scale linked-read sequencing to model the strength and timing of selective sweeps on loci identified to contribute to variation of this carotenoid-based trait.

## Results

Evaluation of Bill Color Variation. We first analyzed bill color variation using UV-vis reflectance spectrophotometry from 948 adult wild-caught and 550 captive-bred long-tailed finches at Macquarie University, Sydney, Australia. Bill color variation is geographically structured across the range of the long-tailed finch into three distinct regions: phenotypically *acuticauda*-yellow populations to the west of the Western Australia – Northern Territory border (~129.1° E), phenotypically *hecki*-red populations to the east of the town of Katherine, Northern Territory (14.5° S, 132.3° E); and phenotypically admixed populations in between (Higgins 2006; Griffith and Hooper 2017; Hooper et al. 2019). We quantified bill hue using the colorimetric variable H3, which is the midpoint between the minimum and maximum reflectance of a surface between 400 and 700 nm (λ_R50;_ Maia et al., 2019). This colorimetric variable efficiently differentiated the yellow and red bills of the two subspecies in our common garden conditions by 56.5 nm (H3: *acuticauda*: 526.1 ± 5 nm [mean ± standard deviation], N = 151; *hecki*: 582.6 ± 6 nm, N = 173; Fig. 1C).

To discern broadscale genetic dominance between long-tailed finch subspecies we next evaluated bill color variation in captive-crossed first-generation (F_1_) hybrids. As a group, F_1_ hybrids had substantially redder bills than might be expected if the allelic contribution of each subspecies was entirely additive (all F_1_ hybrids: 567.2 ± 8 nm, N = 226; hypothetical intermediate: 554.4 nm). In birds, females are the heterogametic sex (i.e., ZW) and males the homogametic sex (i.e., ZZ) so if a recessive Z-linked effect on bill color comes from one subspecies, F_1_ females with that subspecies father should differ from all other hybrids. F_1_ hybrid ZW females with an *acuticauda* father had significantly yellower bills (561.1 ± 7 nm, N = 66) than F_1_ hybrid females with a *hecki* father (571.3 ± 7 nm, N = 58) and both groups of F_1_ hybrid males (ZZ; *acuticauda* father: 567.5 ± 7 nm, N = 51; *hecki* father: 570.0 ± 7 nm, N = 51; *P* < 0.005 for all three comparisons; Fig. 1D). This confirms a recessive sex-linked contribution to bill color, first noted by McDiarmid et al. (2023). There was no significant difference in bill color between F_1_ hybrid females with a *hecki* father and either group of F_1_ hybrid males. The observed interaction between hybrid cross direction and sex strongly suggests that the overall Z-linked allelic contribution from yellow-billed *acuticauda* is recessive to that of red-billed *hecki* (Fig. 1D).

We expanded the analysis of variation in bill hue across the geographic ranges of both subspecies, and the hybrid zone between them, using 948 wild-caught samples (Fig. 1E). Our sampling includes phenotypically “pure” populations from each subspecies at either end of the transect (i.e., >80% of member individuals within two standard deviations of the mean bill hue of each subspecies measured in captive common garden conditions) and a set of populations in between that span nearly the full range of potential color variation between them (Fig. 1F). Only two populations sampled in the wild (pops. 24 and 25) exhibited a mean bill color like that observed in our captive crossed F_1_ hybrids (Fig. 1D and 1F). This suggests that recombination between the allelic variation underlying bill color has been substantial. We leverage this naturally occurring phenotypic variation to identify the underlying genetic basis.

### Each Subspecies has a Distinct Carotenoid Composition in the Bill

We examined the carotenoid composition in the bills of each long-tailed finch subspecies using high-performance liquid chromatography (HPLC) which revealed highly distinctive differences between the two (Fig. 2; Fig S1). We detected five primary carotenoid pigments in the yellow bills of subspecies *acuticauda*: three dietary yellow-orange carotenoids (lutein, zeaxanthin, and β-cryptoxanthin), a metabolized yellow carotenoid (anhydrolutein), and a dietary red carotenoid (lycopene). The red bills of subspecies *hecki* were predominantly comprised of six carotenoid pigments: a dietary yellow carotenoid (lutein), a metabolized yellow carotenoid (anhydrolutein), a dietary red carotenoid (lycopene), and three metabolized red carotenoids (astaxanthin, α-doradexanthin, and adonirubin). Notably, the three dietary yellow-orange carotenoids that occur at highest abundance in *acuticauda* bills are each the direct antecedent of one of the three metabolized red ketocarotenoids found at highest abundance in *hecki* bills (Fig. 2C). Moreover, each of the ketocarotenoids present in the bills of *hecki* are a byproduct of the same metabolic reaction in the form of C(4)-oxidation, a process in which a carbonyl group is added to the C(4) position of a β-ring end group (LaFountain et al. 2015). A set of ε,ε-carotenoids, canary xanthophylls a and b, were detected in low concentrations in yellow bills (Fig. S1). This low concentration is likely to be an artefact, the result of a necessary saponification step used during carotenoid pigment extraction (Toomey et al. 2022b). We therefore refrain from formally comparing total carotenoid content in yellow bills to that in red bills. Altogether, these observations identify C(4)-oxidation as the enzymatic process responsible for the difference in bill color between long-tailed finch subspecies: specifically, a lack of C(4)-oxidation in the yellow bills of *acuticauda*.

**Fig. 2.**
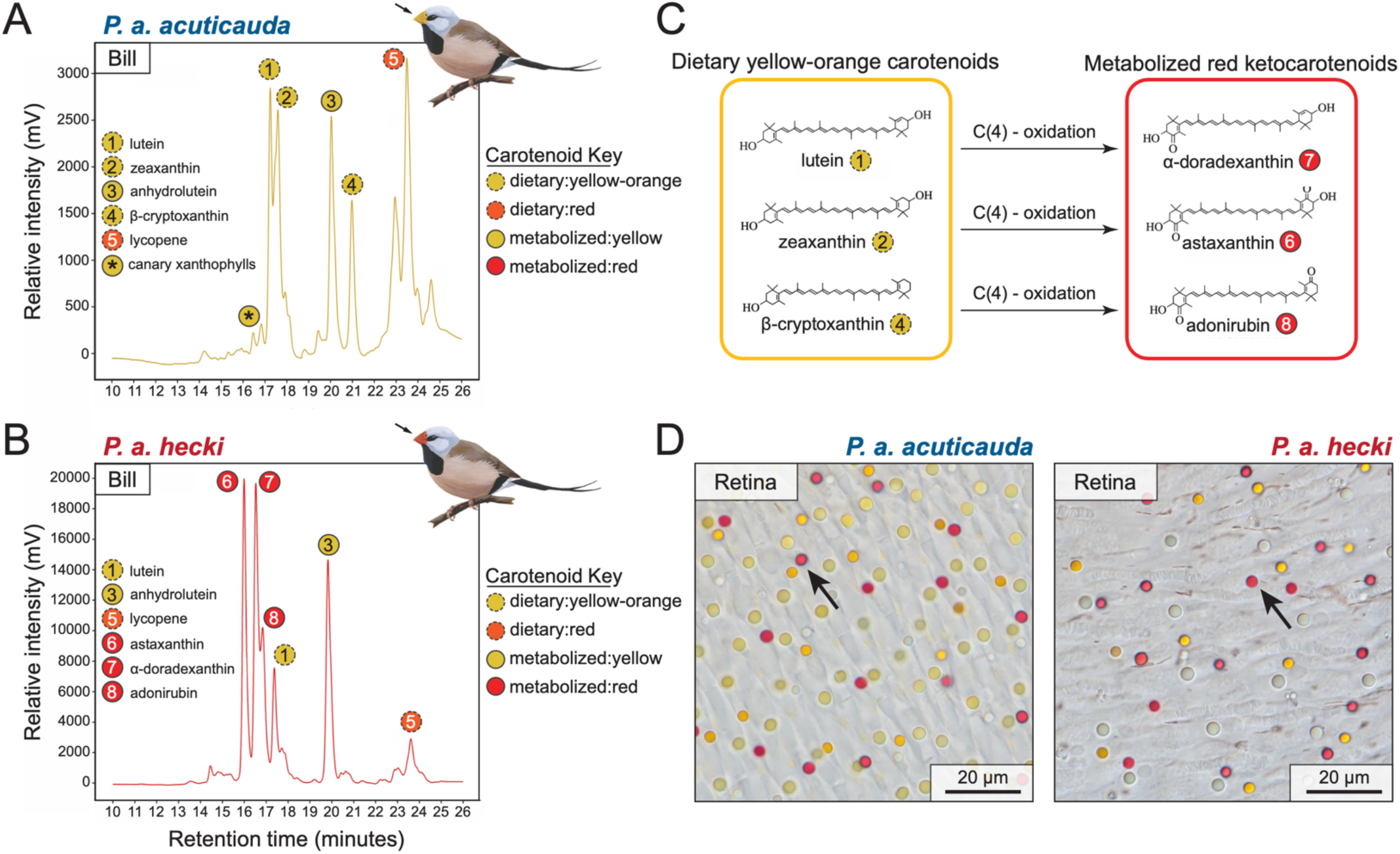
Carotenoid composition of long-tailed finch bills and retinal cone oil droplets. (A and B) High performance liquid chromatography chromatogram for carotenoids isolated from bill tissue integument from individuals of subspecies *acuticauda* (A) and *hecki* (B). Carotenoids were identified via comparison with known standards. Each unique carotenoid has been numerically annotated from left to right first in *acuticauda* and then in *hecki* with each carotenoid labeled as dietary yellow-orange (yellow dashed circle), dietary red (red dashed circle), metabolized yellow (solid yellow circle), or metabolized red (solid red circle). The bills of subspecies *acuticauda* also contained a set of metabolized ε,ε-carotenoids not found in the bills of subspecies *hecki*: canary xanthophylls a and b, which are annotated with an * within a solid yellow circle in panel A. Canary xanthophylls are known to be artificially depleted by the saponification step used during pigment extraction. (C) Metabolic conversions utilized by birds to produce the three red ketocarotenoids found in *hecki* bills from the three dietary yellow-orange carotenoids observed in *acuticauda* bills. All three biosynthetic pathways utilize C(4)-oxidation. (D) Retinal wholemounts of the two subspecies of long-tailed finch: *acuticauda*, on left, and *hecki*, on right. Red cone photoreceptor oil droplets of both subspecies contain the ketocarotenoid astaxanthin, a metabolic byproduct from C(4)-oxidation of the dietary carotenoid zeaxanthin.

### Both Subspecies Synthesize Red Ketocarotenoids in the Retina

We used brightfield microscopy of long-tailed finch retinas to examine the composition of retinal cone photoreceptor cell types in each subspecies and observed no difference between them (Fig. 2D). Most birds possess six subtypes of cone photoreceptor and five of these contain oil droplets with distinctive autofluorescence signatures due to their carotenoid composition (Toomey et al. 2015). The sixth subtype may or may not contain carotenoids, depending on species and retinal location (Hart et al., 1998). We classified cone cell subtypes to identify differences, if any, between the two subspecies in the occurrence of red oil droplets in long-wavelength-sensitive red cone cells. These oil droplets contain the ketocarotenoid astaxanthin (Goldsmith et al., 1984; Toomey et al., 2015), a metabolized byproduct of the dietary carotenoid zeaxanthin (LaFountain et al., 2015; Fig. 2C). Strikingly, the retinas of *acuticauda* show no difference from those of *hecki* in the occurrence of red oil droplet-containing single cone cells (Fig. 2D). This contrasts with the case of the orange-mutant red-throated parrotfinches *Erythrura* psittacea, which are constitutively unable to metabolize red ketocarotenoids, both in the feathers and in retinal cone cells (Toomey et al. 2022a). Our results strongly suggest that the genetic basis underlying the principal color difference between long-tailed finch subspecies is regulatory as *acuticauda* can utilize C(4)-oxidation to convert dietary carotenoids into metabolized red ketocarotenoids within the retina even though they do not do so in the bill integument.

### Genomic Differentiation

We used the haplotagging approach described in Meier et al. (2021) to generate whole genome linked-read (LR) sequence data for the long-tailed finch and its closely related allopatric sister species the black-throated finch *P. cincta* (diverged 1.8 million years ago; Lopez et al. 2021). We sequenced 1133 *P. acuticauda* (both subspecies) and 96 *P. cincta* samples in 96-plex batches to a median read coverage of 1.38× with samples both individually and molecularly barcoded; see *Materials and Methods*). Across the 1204 samples that had high molecular weight DNA available for haplotagging, we recovered a mean molecule N50 of 12.2 kbp (± 4.1 kbp) with maximum molecule sizes averaging 106.3 kbp (± 22.1 kbp; Fig. S2). Following variant calling and imputation, we retained a set of 29.3 million SNPs and observed 3.9 million fixed differences in both *Poephila* species relative to the zebra finch reference genome.

Background genomic differentiation between long-tailed finch subspecies is highly skewed onto the Z chromosome (mean genetic distance F_ST_ for autosomes: 0.027; for chrZ: 0.551; Fig. 3A). This observation is consistent with a previous finding that differentiation is likely associated with a large Z-linked inversion that acts as a barrier to gene flow between subspecies (Hooper et al., 2019). We used a set of 649 linkage disequilibrium (LD) pruned ancestry informative markers (defined as SNPs with an allele frequency (AF) difference > 0.8 between allopatric pops. 1 – 7 and pops. 28 – 34) to calculate a hybrid index between subspecies. We estimated the hybrid zone to be 126.9 km wide (107.8 – 148.7 km, 95% highest posterior density interval) along the edge of the Kimberley Plateau, Western Australia (between populations 8 and 19, see Fig. 1E and Fig. S3). Remarkably, individuals from hybrid zone populations exhibit bill color that is on average just as yellow as individuals from pure *acuticauda* populations (H3: *acuticauda* 535.2 ± 10 nm, N = 265; hybrid zone 534.7 ± 10 nm, N = 158; *P* = 0.9, Tukey’s HSD test). As a result, the long-tailed finch hybrid zone is effectively cryptic with respect to bill color. As *hecki* alleles appear in aggregate to be dominant (Fig. 1D), this would suggest that the *acuticauda* alleles for yellow bill coloration have asymmetrically introgressed into *hecki* following secondary contact, resulting in populations featuring yellow-billed *hecki* birds.

**Fig. 3.**
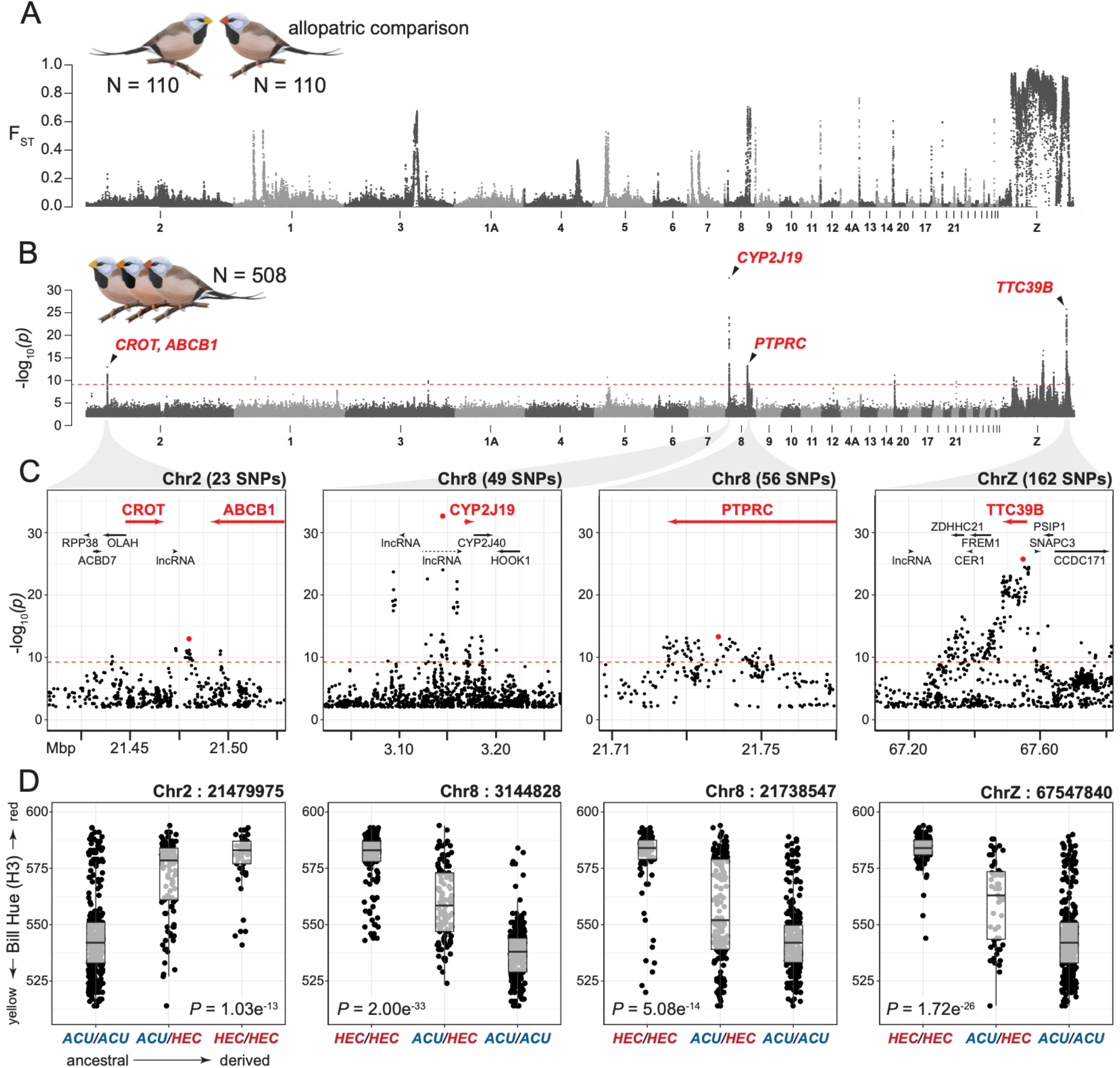
Genome wide association studies identify four regions underlying most variation of bill color in the long-tailed finch. (A) Genomic differentiation (F_ST_) between allopatric *acuticauda* (pops. 1-7; N = 110) and allopatric *hecki* (pops. 28-34; N = 110) is concentrated on the Z chromosome. F_ST_ was calculated after excluding singleton sites in 20 kb windows with 10 kb step size. Chromosomes are ordered from left to right as the largest to smallest autosome and then the Z chromosome. Only male samples were used to calculate F_ST_ on the Z chromosome (*acuticauda*: N = 70; *hecki*: N = 69) to circumvent any potential issues with hemizygosity in females. (B) Manhattan plot for GWAS of bill color hue (H3) from 508 long-tailed finches identified eleven association peaks. Association support plotted as the −log10(Pwald). Dashed red line denotes the genome-wide significance threshold. Genes with carotenoid processing function are annotated in red for the top four association peaks. (C) Zoom-in to the top four association peaks in (B). Genes within each window are annotated and those linked with carotenoid processing shown in red. The most strongly associated SNP in each window is indicated with a red circle. (D) Genotype by phenotype (H3) boxplots for the most strongly associated SNP in each window in (C). Genotypes are given with alleles polarized ancestral or derived from left to right with bill color hue (H3) given in nanometers on the y-axis. The GWAS p-value is given for each SNP as an inset.

### Association Mapping

We performed genome-wide association studies (GWAS) to identify loci underlying bill hue while controlling for population structure by including an inter-individual relatedness matrix, hybrid index scores, and sex as covariates. We found eleven association peaks that together explained 93.8% of the variance in this trait (Table 1). Four of these association peaks together account for 53% of variance (Fig. 3B). We describe each of these in order of descending amount of variance explained.

**Table 1.** Summary of eleven bill color association peaks identified by genome-wide association studies. Association peak window size determined based on the location of the first and last SNP above genome-wide significance separated by less than 50 kbp. Candidate genes in bold are those which are discussed in this manuscript. F_ST_ is reported between allopatric populations of each subspecies. Results from GEMMA analysis are provided for the most-significant SNP in each window.

| Chrom. | Window (Mb) | Window size (kbp) | SNPs | Candidate Genes | Top SNP | F<sub>ST</sub> | AF | $\beta$ | P-value | Genetic Variance |
| --- | --- | --- | --- | --- | --- | --- | --- | --- | --- | --- |
| chr2 | 21.44 - 21.50 | 55.43 | 23 | **CROT / ABCB1** | chr2:21479975 | 0.419 | 0.219 | 8.09 | 1.03e<sup>-13</sup> | 0.061 |
| chr8 | 3.09 - 3.20 | 112.37 | 54 | **CYP2J19** | chr8:3144828 | 0.952 | 0.402 | 11.99 | 2.00e<sup>-33</sup> | 0.216 |
| chr8 | 21.72 - 21.75 | 29.02 | 57 | **PTPRC** | chr8:21738547 | 0.547 | 0.336 | 7.31 | 5.08e<sup>-14</sup> | 0.065 |
| chr20 | 0.32 - 0.47 | 153.16 | 5 | *RBM39 / PHF20* | chr20:365722 | 0.786 | 0.336 | 6.91 | 7.77e<sup>-12</sup> | 0.057 |
| chrZ | 13.32 - 13.48 | 167.95 | 15 | *RORB* | chrZ:13402137 | 0.176 | 0.342 | -6.37 | 1.89e<sup>-11</sup> | 0.059 |
| chrZ | 42.99 - 43.77 | 782.32 | 80 | *SLC45A2* | chrZ:43673992 | 0.015 | 0.240 | -8.6 | 2.56e<sup>-17</sup> | 0.086 |
| chrZ | 54.11 - 54.46 | 346.60 | 42 | *MAST4 / CD180* | chrZ:54299700 | 0.058 | 0.267 | -6.86 | 1.79e<sup>-12</sup> | 0.058 |
| chrZ | 67.29 - 67.62 | 329.61 | 207 | **TTC39B** | chrZ:67547840 | 1.000 | 0.273 | 11.57 | 1.72e<sup>-26</sup> | 0.189 |
| chrZ | 67.73 - 67.87 | 144.57 | 17 | *CCDC171* | chrZ:67725101 | 0.884 | 0.316 | 7.36 | 4.60e<sup>-13</sup> | 0.017 |
| chrZ | 68.07 - 68.22 | 146.76 | 23 | *BNC2* | chrZ:68132346 | 0.103 | 0.327 | -7.52 | 6.48e<sup>-15</sup> | 0.082 |
| chrZ | 70.12 - 70.40 | 278.26 | 39 |  | chrZ:70241903 | 0.020 | 0.210 | -6.86 | 1.67e<sup>-11</sup> | 0.048 |

The most strongly associated peak, located on chromosome 8, is composed of 54 SNPs with association values above the genome-wide significance threshold. This peak spans three protein-coding genes, including the oxidative ketolation gene *CYP2J19* (Fig. 3C and Table 1). The SNP most significantly associated with bill hue (chr8:3144828, P = 2.00e^−33^; Fig. 3D) explains 21.6% of variance (*β* = 12.0 nm) and is located 21.9 kbp upstream of *CYP2J19* within a long non-coding RNA (lncRNA). This variant, and another 420 bp away, are the only two autosomal SNPs with F_ST_ > 0.95 between allopatric populations of each subspecies. We found 10 SNPs above genome-wide significance located within the genic domain of *CYP2J19*, all within introns, and a further 13 SNPs located less than 20 kbp upstream. Previous laboratory-based studies revealed that *CYP2J19* – together with *BDH1L* – is an essential component of ketocarotenoid production in birds and other vertebrates (Lopes et al., 2016; Mundy et al., 2016; Toomey et al., 2022a). Recent admixture mapping efforts in other natural avian systems have also found an association between *CYP2J19* and carotenoid-based color variation (Kirschel et al., 2020; Aguillon et al., 2021; Khalil et al., 2023).

The second most strongly associated peak, located on the Z chromosome, was composed of 207 SNPs above our genome-wide significance threshold and contained seven protein-coding genes, including the oxidative ketolation enhancer gene *TTC39B* (Fig. 3C and Table 1). The most significant SNP in this association peak (chrZ:67547840, P = 1.72e^−26^; Fig. 3D) is the second most significant SNP genome-wide and explains 18.9% of variance (*β* = 11.6 nm). The chrZ:67547840 SNP is located within the first intron of *TTC39B* and is fixed (i.e., F_ST_ = 1.0) between long-tailed finch subspecies. A total of 53 SNPs above genome-wide significance were located within the genic domain of *TTC39B*, seven of which are 3’ UTR variants and one of which is a synonymous substitution. An additional 12 SNPs above genome-wide significance were located less than 20 kbp upstream of the gene. When co-expressed with *CYP2J19*, *TTC39B* greatly enhances the conversion of yellow dietary carotenoids to red ketocarotenoids (Toomey et al., 2022a) and has been shown from admixture studies to be significantly associated with yellow-red color variation in both birds and fish (Hooper et al., 2019; Ahi et al., 2020; Toomey et al., 2022a).

A third association peak, also located on chromosome 8, includes 57 SNPs above genome-wide significance, all of which are located within the genic domain of the protein tyrosine phosphatase receptor type C gene *PTPRC* – also known as *CD45* (Fig. 3C and Table 1). This large transmembrane glycoprotein, which has six isoforms in humans (Hermiston et al. 2003) and seven in the zebra finch (Rhie et al., 2021), is found on the cell surface of all hematopoietic cells other than mature erythrocytes and plays a role in innate immune response across different cell types (Al Barashdi et al. 2021). The most significant SNP in this association peak (chr8:21738547, P = 5.08e^−14^; Fig. 3C) explains 6.5% of variance (*β* = 7.3 nm). Eleven of the associated SNPs are located within exons, but only two encode non-synonymous substitutions (R401G and A864G), two are synonymous substitutions, and seven are in the 3’ UTR. While a prior study of the long-tailed finch also found SNPs near *PTPRC* to be associated with bill color variation (Hooper et al., 2019), there have been no other direct links with carotenoids or color variation in other systems. One possible causal association between *PTPRC* and bill color comes from the fact that carotenoids – especially astaxanthin – have well-established immune system functions in humans due to their potent antioxidant and anti-inflammatory properties (Hussein et al., 2006; Fakhri et al., 2018), although the role of carotenoids in immune function of birds has been challenged (Koch et al., 2018).

The fourth peak, located on chromosome 2, spans 23 SNPs and contains five protein-coding genes (Fig. 3C and Table 1). One of these genes, *ABCB1*, is an ATP-binding cassette (ABC) transporter that translocates phospholipids across cell membranes. Members of its gene family (*ABCA1* and *ABCG1*) have been associated with the deposition of carotenoids in the retina of chickens (Connor et al., 2007) and the red feathers of red-backed fairywrens (Khalil et al., 2022), respectively. The most significant SNP in this association peak (chr2: 21479975, P = 1.03e^−13^; Fig. 3D) explains 6.1% of variance (β = 8.1 nm) and is located 11.7 kbp downstream of ABCB1 and three SNPs above genome-wide significance are located within its third intron. Two other genes of interest in this association peak, *OLAH* and *CROT*, are involved in the release of free fatty acids from fatty acid synthetase (*FASN*) and mitochondrial fatty acid β-oxidation, respectively. Two SNPs above genome-wide significance are located within an intron of *OLAH*. While these two genes have no prior association with carotenoids or color variation, their role in lipid metabolism within the mitochondria is notable as recent evidence suggests that carotenoid ketolation and ornamentation is functionally linked to mitochondrial performance (Cantarero and Alonso-Alvarez 2017; Cantarero et al., 2020; Hill et al., 2019). More specifically, the ketolation of dietary carotenoids to red ketocarotenoids occurs through the addition of a ketone (or carbonyl) group. Ketones are a metabolic byproduct of the mitochondrial fatty acid β-oxidation pathway that *CROT* belongs to and as such may constitute a target for modulating ketocarotenoid production (Houten and Wanders 2010).

In summary, we find variation in bill hue in the long-tailed finch is associated most strongly with regions of the genome that include genes known to be involved in – or are plausibly linked with – carotenoid processing. The top two association peaks contain *CYP2J19* and *TTC39B*. *CYP2J19* is one of two essential enzymes for the conversion of dietary carotenoids to red ketocarotenoids via C(4)-oxidation and *TTC39B* enhances carotenoid metabolism, respectively (Toomey et al., 2022a). Each of the four top association peaks contained a locus of major effect (i.e., PVE > 5%). We used outgroup information from the black-throated finch and zebra finch to polarize alleles associated with bill color variation to their subspecies of origin. We found that the *acuticauda* allele is the derived condition at the most significant SNP in three of the four top association peaks – regions including *CYP2J19, PTRPRC*, and *TTC39B* (Fig. 3D). In contrast, while the derived allele at the most significant SNP on chromosome 2 is common in *hecki* (AF = 0.41) it is also found at similar frequency in the black-throated finch (AF = 0.36). This suggests that the yellow bill color of *acuticauda* is the derived phenotypic condition in the long-tailed finch.

### Geographic and Genomic Clines Support Biased Introgression of Bill Color Alleles

Focusing on yellow billed variants, we observe clear evidence of introgression at *CYP2J19* and *TTC39B* from yellow-billed *acuticauda* into red-billed *hecki*. The geographic cline centers for the two SNPs most strongly associated with bill color variation are shifted 220.4 km (*CYP2J19*: chr8:3144828, center = 868.0 km from most western sampled population [849.7 – 885.3 km 95% HPDI]) and 373.5 km (*TTC39B*: chrZ:67547840, center = 1021 km [993.7 – 1051.0 km 95% HPDI]) to the east of the center of genomic admixture between subspecies (hybrid index, center = 647.6 km [642.4 – 652.8 km 95% HPDI]), respectively (Figure 4). While the introgressing alleles at both loci originated within subspecies *acuticauda*, they differ in notable aspects of their respective clines and underlying allele frequencies in color-admixed populations (i.e., populations 20 – 27 located between dashed lines 1 and 2 in Figure 4A-C). First, the cline center for *TTC39B* (chrZ:67547840) is located a further 153 km east of the center for *CYP2J19* (chr8:3144828) (Figure 4D). Second, the *acuticauda CYP2J19* allele at SNP chr8:3144828 was observed segregating at intermediate frequency within color-admixed populations: allele frequencies ranging from 0.65 in the west [pop. 20] to 0.16 less than 80 km to the east [pop. 26] (Figure 4B). In contrast, the *acuticauda TTC39B* allele at SNP chrZ:67547840 was observed as fixed or nearly fixed in these same populations (ranging from 1.00 to 0.86; Figure 4C). The *acuticauda* allele of SNP chrZ:67547840 was never observed at a frequency less than 0.86 in populations where both alleles were present.

**Fig. 4.**
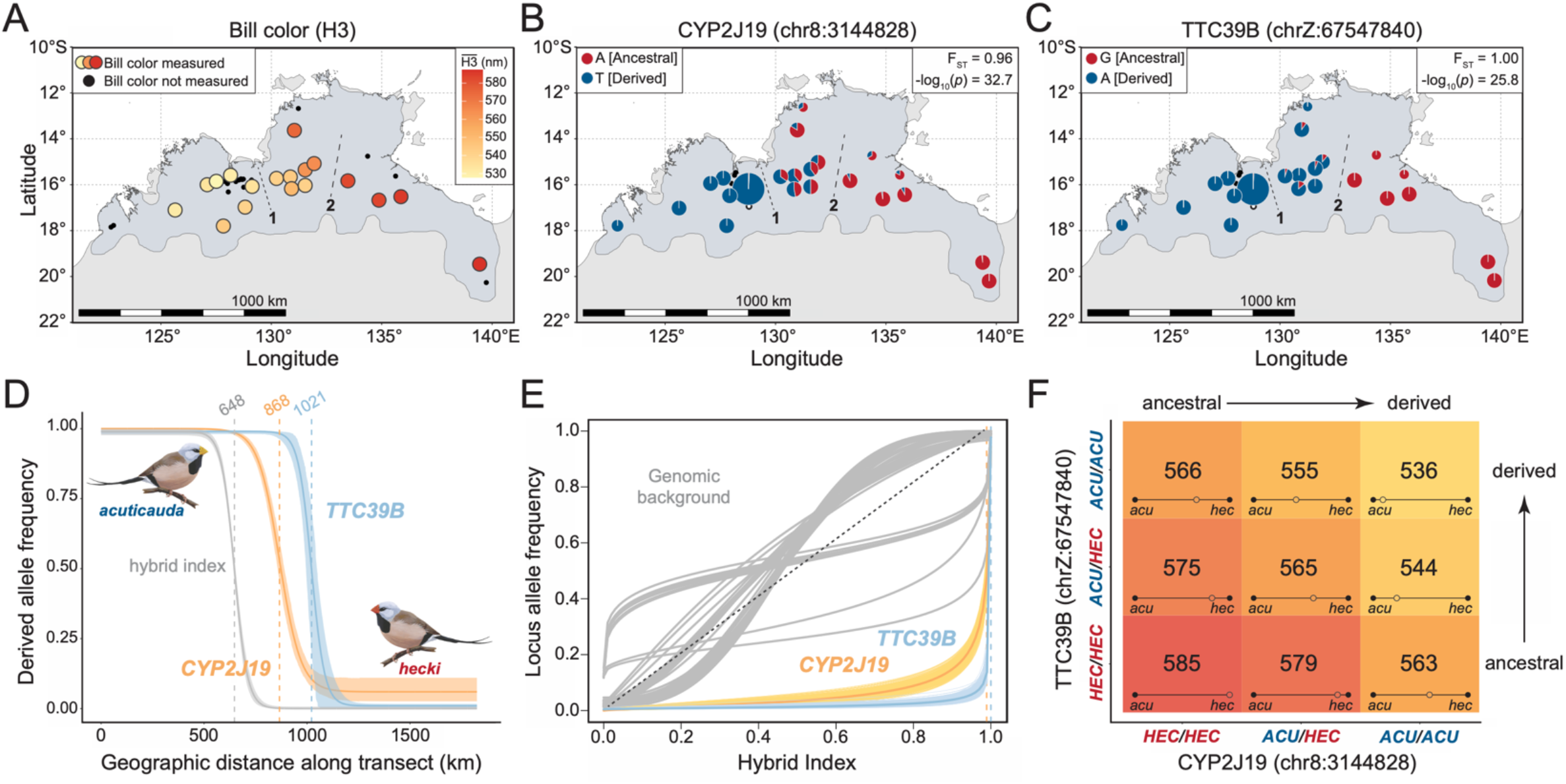
Evidence from geographic and genomic clines support an epistatic relationship between *CYP2J19* and *TTC39B*. (A) Geographic variation in bill hue (H3) across the 18 populations of the long-tailed finch where bill color was measured. Populations between dashed lines 1 and 2 represent color-admixed *hecki* (pops. 20-27) subject to introgression from subspecies *acuticauda*. Immediately west of dashed line 1 are yellow-billed populations from the genomic hybrid zone between subspecies. Immediately east of dashed line 2 are populations from red-billed *hecki* (pops. 28-34). (B and C) Population allele frequencies for the two SNPs most strongly associated with color variation. Derived and ancestral alleles are color-coded blue and red, respectively. F_ST_ and GWAS significance given in top right inset of each panel. (D) Geographic clines for genome-wide hybrid index (grey cline), *CYP2J19* (chr8:3144828, orange cline), and *TTC39B* (chrZ:67547840, blue cline). Best fit clines shown as solid clines with 95% HPDI shading. Vertical dashed lines with values in kilometers atop represent inferred cline centers. (E) Genomic cline analysis indicates that introgression of variants associated with *CYP2J19* (chr8:3144828, orange) and *TTC39B* (chrZ:67498406, blue) has been significant relative to genomic background (grey). Genomic background represented as genomic clines for 100 randomly drawn ancestry-informative markers used to infer hybrid index. (F) Allelic dominance and evidence of epistasis between *CYP2J19* and *TTC39B*. Each grid cell represents a genotype combination observed in long-tailed finches at the two SNPs most strongly associated with color variation. The mean bill hue (H3) for each genotype combination is given within each grid cell in nanometers (nm). The phenotypic position of each genotype combination is shown along the 55 nm color gradient between yellow-billed *acuticauda* (mean H3 = 530 nm, N = 243, pops. 2-10) and red-billed *hecki* (mean H3 = 585 nm, N = 185, pops. 28-33).

Genomic cline analysis bolsters the significance of introgression of *acuticauda* yellow alleles at bill color genes *CYP2J19* and *TTC39B* (Figure 4E). Of the 649 LD-pruned ancestry informative markers (see Materials and Methods) used to calculate a hybrid index between long-tailed finch subspecies, the three SNPs with the strongest support for introgression were all located within 38 kbp of *CYP2J19* and the most significant was the top GWAS SNP chr8:3144828 (*c* = 0.994, P = 5.90e^−201^). The SNP with the fourth strongest support for introgression was located within an intron of *TTC39B* (chrZ:67498406, *c* = 1, P = 2.14e^−71^) and was also significantly associated with bill color variation (GWAS P = 5.92e^−23^). Due to LD-pruning, the *TTC39B* SNP used in geographic cline analysis (chrZ:67547840) was not included in the set of ancestry informative markers used for genomic cline analysis. Instead, another tightly linked SNP, chrZ:67498406 (49.4 kbp away; r^2^ = 0.90, D’ = 0.95), is among the set of ancestry informative markers. Three SNPs on chromosome 20 (Table 1) exhibited evidence of introgression from *acuticauda* into *hecki*, one of which was also significantly associated with bill hue variation (chr20:317305, *c* = 0.997, P = 1.48e^−56^; GWAS P = 4.1e^−10^). This SNP is located 1.2 kbp upstream of RNA-binding protein 39 (*RBM39*).

Taken together, results of geographic and genomic cline analyses showcase the introgression of *acuticauda* alleles for yellow bill coloration into a genomic background that is otherwise that of red-billed *hecki* (Figure 4E). The ~150 km separation between the geographic cline centers for these two loci suggest in turn that each allele is introgressing under distinct evolutionary regimes (Figure 4D). We did not perform clinal analyses within any other association peak region (Table 1) because no SNPs within these were identified as ancestry informative (i.e., F_ST_ > 0.7 and ΔAF > 0.80 between allopatric populations). Without speculating on the exact mechanism under which yellow bill color may have been favored by selection, the evidence nonetheless suggests that yellow bill color, and/or a tightly linked trait, is conferring a selective advantage that has promoted their asymmetrical introgression from *acuticauda* into *hecki*.

### Evidence of Epistasis Between *CYP2J19* and *TTC39B*

A biochemical relationship between the products of *CYP2J19* and *TTC39B* has been established experimentally, which identified the latter as a potent enhancer of ketocarotenoid biosynthesis (Toomey et al., 2022a). Toomey et al. (2022a) proposed that *TTC39B* acts to facilitate the transport of carotenoids to or from the location of enzymatic conversion. We leveraged the difference in cline centers to explore epistatic effect on phenotype, if any, between *CYP2J19* and *TTC39B* in a natural system. To do so, we quantified mean bill hue (H3) using the 508 individuals with phenotype data carrying zero, one, or two copies of the *acuticauda* allele at chr8:3144828 (*CYP2J19*) and chrZ:67547840 (*TTC39B*). We found that the red *hecki* allele is dominant at *CYP2J19* and that *TTC39B* alleles appear to be additive in their effect (Figure 4F). Individuals carrying a single copy of the red *hecki* allele at *CYP2J19* were indistinguishable in hue from those carrying two copies if they were also homozygous for the *hecki* allele at *TTC39B* (homozygous *hecki*, H3 = 584.6, N = 99; heterozygous *hecki*, H3 = 579.4, N = 13, Figure 4F). Consistent with an epistatic effect on phenotype between *CYP2J19* and *TTC39B*, individuals carrying at least one copy of the *hecki* allele at *CYP2J19* exhibited an increase in bill hue of 10 to 15 nm for each copy of the *hecki* allele they carried at *TTC39B* (Figure 4F). The recessive nature of the yellow *acuticauda* allele of *CYP2J19* is consistent with the role of this enzyme as an oxidative ketolase while the additive contribution of each *TTC39B* is consistent with its shown role as an enhancer of carotenoid metabolism (Toomey et al., 2022a). The importance of considering the effect on bill hue of these two loci together is made further evident by contrast with the seemingly additive contribution of each *CYP2J19* allele when this locus is considered in isolation (see Figure 3D).

Results from naturally admixed birds are largely consistent with the inferences drawn from bill color variation observed in captive crossed F_1_ hybrids: namely that red *hecki* alleles are net dominant to yellow *acuticauda* alleles (Figure 1D). While the net contribution of *acuticauda* alleles on the Z chromosome appears to be recessive in captive-crossed F_1_ hybrids (Figure 1D), the *acuticauda* allele of Z-linked *TTC39B* appears to be additive in admixed birds in the wild (Figure 4F). This difference is likely to represent the combined contribution of additional autosomal and Z-linked loci of smaller effect on bill color variation we identified (Figure 3B and Table 1).

### Signatures of Selection at Bill Color Associated Genes

The initial divergence in bill color between long-tailed finch subspecies and more recent introgression of bill color alleles from *acuticauda* into *hecki* may have been driven by an adaptive benefit for individuals with yellower bills. To explore this, we scanned for signatures of selective sweeps within each of the four regions most strongly associated with bill color variation in haplotype homozygosity summary statistics and by reconstructing local gene trees, to approximate the ancestral recombination graph (ARG). Selective sweeps are expected to result in an increase in haplotype homozygosity around a target of selection (Vitti et al., 2013) and a decrease in the time to coalescence for haplotypes carrying a favored allele (Hejase et al. 2020; Stern et al. 2019). We focus on ARG-based inference as these approaches are more informative and direct compared with haplotype homozygosity statistics and site frequency spectrum (SFS) approaches; which are themselves low-dimensional summaries of the ARG (Vitti et al. 2013; Speidel et al., 2019; Shipilina et al. 2022).

We phased all variants on chromosomes 2, 8, and Z in samples with linked-read information using HapCUT2 (Edge et al. 2017). Linked-read (LR) information greatly increased phasing performance: we recovered an 18-fold improvement in the length of phased blocks, with a median N50 of 19.6 kbp using LR data (versus 1.1 kbp for short-read (SR) data; Fig. S4). Of the 18 samples used as technical replicates, effectively controlling for variation in high-molecular weight DNA quality, phased block N50s were on average 29-fold longer when utilizing LR information (LR: 20.4 kbp, SR: 0.7 kbp) despite these samples having an average depth of coverage 13-fold lower (LR: 2× depth, SR: 26×; Fig. S4). We next used this LR-phased haplotype data to generate ARGs from our four top association peaks ± 1 Mb using Relate v1.1 (Speidel et al. 2019) and screened for lineages carrying mutations that have spread faster than competing lineages. As the Relate Selection Test assumes no population stratification (Speidel et al. 2019), we evaluated support for selective sweeps in *acuticauda* (pops. 1 – 7) and *hecki* (pops. 20 – 34) separately and did not test for selection within the hybrid zone populations between them.

We found evidence of selective sweeps on SNPs associated with bill color variation in the long-tailed finch. Within the genomic window on chromosome 8 containing *CYP2J19*, the *acuticauda* derived variant at bill color associated SNP chr8:3094115 (GWAS P = 5.45e^−19^) exhibited evidence of strong selection in subspecies *hecki* (Fig. 5, s = 0.0059, logLR = 34.2). The variant is nearly fixed within *acuticauda* (pops. 1 – 7, AF = 0.99) and the hybrid zone (pops. 8 – 19, AF = 0.99). It appears to be approaching fixation within color-admixed *hecki* (pops. 20 – 27, AF = 0.83) but is currently at much lower frequency in red-billed *hecki* (pops. 28 – 34, AF = 0.21; Fig. 5B). Consistent with introgression following secondary contact, the sweeping variant first appeared in *acuticauda* approximately 40 kya and has subsequently undergone a rapid increase in frequency within *hecki* between 5 kya and the present day (Fig. 5C). Haplotype homozygosity statistics offer insights into the complicated history of selection on the genomic region encompassing *CYP2J19*. Consistent with a recent selective sweep, cross-population extended haplotype homozygosity (xpEHH) is significantly greater in color-admixed compared to red-billed populations of hecki at SNP chr8:3096231 (-log10(P) = 13.4; Fig. S9); a site only 2 kbp away from the SNP identified via ARG based inference above. The SNP most strongly associated with bill color variation in the long-tailed finch (chr8:3144828; GWAS P = 2.00e^−33^) also had the greatest xpEHH support of any SNP between *acuticauda* and color-admixed populations of *hecki* (-log10(P) = 17.3; Fig. S9). The increase in homozygosity for haplotypes carrying the *acuticauda* derived allele is consistent with a selective sweep at this site, which is located within a lncRNA gene 20 kbp upstream of *CYP2J19*. Extended haplotype homozygosity (EHHS) inference is more consistent with the sweep having occurred within *acuticauda*. Haplotypes carrying the derived variant were on average 3-9× longer in *acuticauda* than in *hecki* (integrated EHH: *acuticauda* = 1596 bp, color-admixed = 472 bp, red-billed = 164 bp; Fig. S13).

**Fig. 5.**
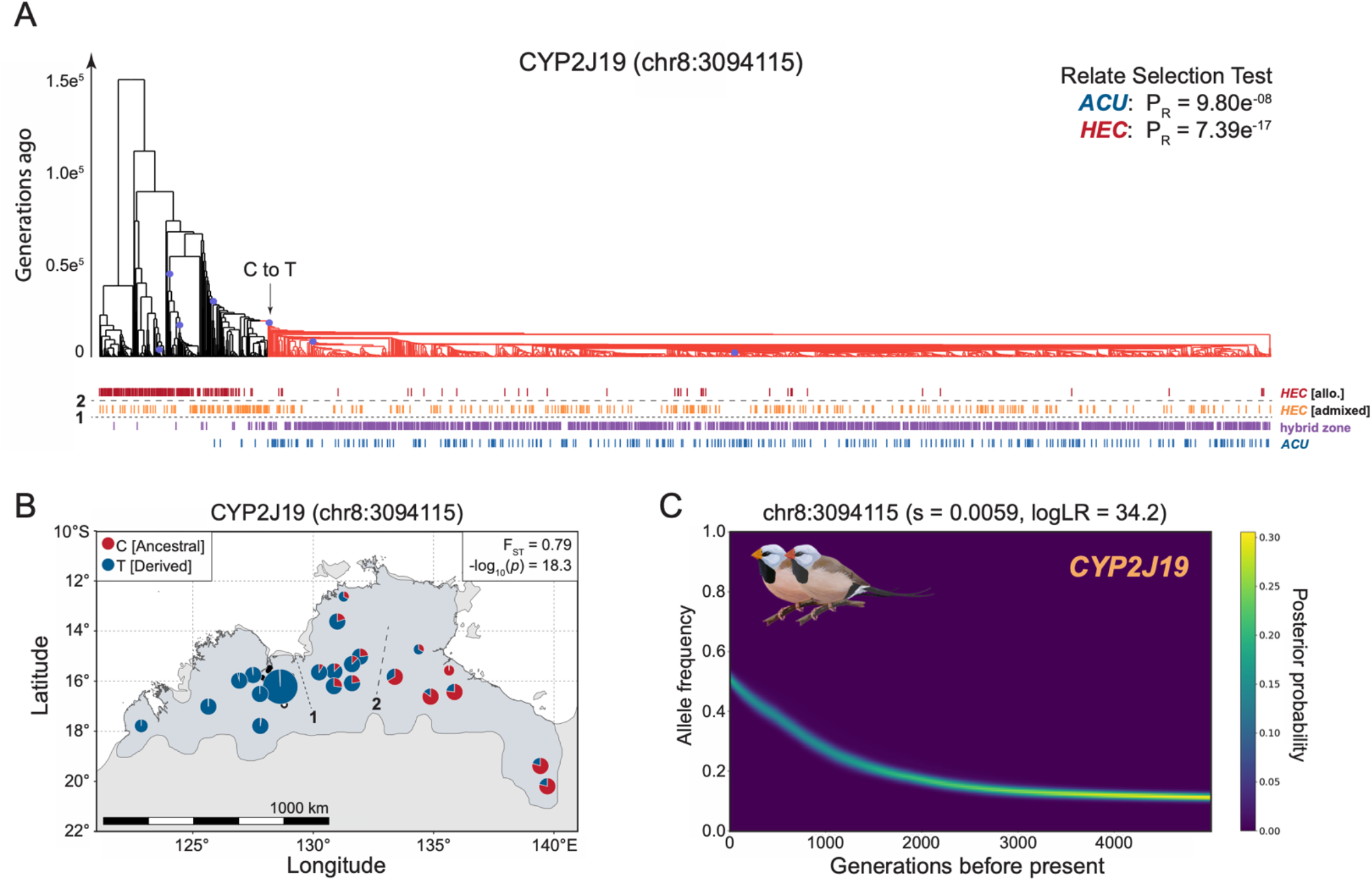
Evidence of selection on *CYP2J19* from ancestral recombination graph (ARG) inference. (A) Relate marginal tree for bill color variation associated SNP chr8:3094115 (2N = 1928 haplotypes). Branches colored in light red represent haplotypes carrying the derived allele at this site. Relate Selection Test evidence for selection over the lifetime of the mutation (P_R_) shown in top right for subspecies *acuticauda* (*ACU*: pops. 1-7, 2N = 282) and subspecies *hecki* (HEC: pops. 20-34, 2N = 522). Vertical hash marks beneath tips of marginal tree represent haplotypes from allopatric red-billed *hecki* (dark red, pops. 28-34), color-admixed hecki (orange, pops. 20-27), the hybrid zone (purple, pops. 8-19), and allopatric yellow-billed *acuticauda* (pops. 1-7). Horizontal dashed lines 1 and 2, as seen in panel (B), represent geographic breaks between the hybrid zone and color-admixed *hecki* (line 1) and between color-admixed and red-billed *hecki* (line 2). (B) Population allele frequencies for SNP chr8:3094115. Derived and ancestral alleles are color-coded blue and red, respectively. F_ST_ and GWAS significance given in top right inset of panel. (C) Historical allele trajectory of the *acuticauda*-derived allele at chr8:3094115 in subspecies *hecki*. A generation time of two years was used to infer timing of selection. Allele trajectories inferred from posterior distribution of marginal trees, inferred with Relate version 1.1 (Speidel et al., 2019), using CLUES (Stern et al., 2019). s, selection coefficient.

Variants within the genomic window on chromosome 8 containing *PTPRC* also exhibited a recent increase in frequency consistent with a selective sweep. An *acuticauda* derived variant at the bill color associated SNP chr8:21738547 (GWAS P = 5.08e^−14^), located within an intron of *PTPRC* and the most significantly associated SNP in this window, shows evidence of strong selection within both subspecies (*acuticauda*, s = 0.0054, logLR = 53.4; *hecki*, s = 0.0076, logLR = 75.3; Fig. S6). This variant is at high frequency in *acuticauda* (AF = 0.78), the hybrid zone (AF = 0.90), and color-admixed *hecki* (AF = 0.83) but is relatively rare in red-billed *hecki* (AF = 0.15; Fig. S6B). The variant appears to have increased in frequency first within *acuticauda* 4-5 kya and has subsequently undergone a rapid ascent in frequency within color-admixed populations of *hecki* between 3 kya and the present day (Fig. S6C). In concurrence with ARG based inference of selection, the SNP with the greatest xpEHH support between color-admixed and red-billed *hecki* populations (chr8:21746905, −log10(P) = 18.1; Fig. S10), is also located within an intron of *PTPRC* and is significantly associated with bill color variation (GWAS P = 2.79e^−10^). Consistent with a recent selective sweep on *acuticauda* derived alleles of *PTPRC*, this SNP exhibits a strong deviation of site-specific extended haplotype homozygosity (EHHS) between color-admixed and red-billed *hecki* populations: 33× longer for carriers of the *acuticauda* derived variant in the former (integrated EHH: *acuticauda* = 5751 bp, color-admixed = 11005 bp, red-billed = 334 bp; Fig. S14).

The genomic window on chromosome 2 containing several candidate carotenoid affiliated genes also exhibited signatures of recent selective sweeps. We observed a sweep-like signature in subspecies *acuticauda* associated with the gene *CROT* centered around SNP chr2:21467694 (s = 0.0071, logLR = 37.6; Fig. S7). The variant showing evidence of a sweep is segregating at intermediate frequency within both *acuticauda* (AF = 0.52) and the hybrid zone (AF = 0.56). It is found at lower frequency within color-admixed *hecki* (AF = 0.14) – increasing in local abundance with proximity to the hybrid zone – and is entirely absent within red-billed *hecki* (AF = 0.00; Fig. S7B). Allele trajectories indicate that this *acuticauda* derived variant has increased in frequency between 4 kya and the present (Fig. S7C). Consistent with this sweep-like signature on *CROT*, a SNP located within intron six (chr2:21456972) had the greatest xpEHH support in *acuticauda* against color-admixed (-log10(P) = 28.4) and red-billed *hecki* (-log10(P) = 15.9; Fig. S11). The derived variant exhibited a strong signature of site-specific extended haplotype homozygosity (EHHS) against *hecki* populations: >12× longer for carriers of the derived variant in *acuticauda* (integrated EHH: *acuticauda* = 3896 bp, color-admixed *hecki* = 283 bp, red-billed *hecki* = 333 bp; Fig. S15).

Compared to autosomal loci, evidence for selective sweeps was more difficult to interpret on the Z chromosome. Despite patterns consistent with selection from geographic and genomic cline analyses (Fig. 4), neither chrZ:67498406 nor chrZ:67547840 shows evidence of selection in *hecki* based on ARG analysis. This appears to be the result of the degree of haplotype divergence between subspecies in this genomic window and the size of the introgressing region (Fig. S8). Indeed, a highly differentiated region approximately 0.4 Mbp in length (from 67.5 to 67.9 Mb, mean allopatric F_ST_ = 0.75), encompassing *TTC39B*, shows evidence of introgression from *acuticauda* into *hecki* (Fig. S12). Ancestral recombination graph approaches for detecting natural selection may be poorly suited in cases of adaptive introgression between highly divergent taxa (Hejase et al., 2020), as such cases violate assumptions regarding the recent coalescence for haplotypes carrying selected alleles, and this problem might be especially pronounced for genomic regions with reduced rates of recombination such as avian sex chromosomes. Haplotype homozygosity statistics also failed to detect any signatures of selective sweeps within this region of the Z chromosome (Fig. S12). However, these methods become power limited as variants approach fixation between focal populations (Stern et al., 2021), as is the case for most variants in this region of the Z chromosome. For example, in populations of color-admixed *hecki*, haplotypes carrying the *acuticauda* derived allele at chrZ:67547840 were nearly fixed (AF = 0.94) and had extended haplotype homozygosity (EHHS) twice the length of those carrying the ancestral allele (ancestral: 8246 bp, derived: 17004 bp) (Fig. S16). Comparison with *acuticauda* and red-billed *hecki* populations is made difficult, however, due to the fixation of alternative alleles at this locus (i.e., F_ST_ = 1.0).

Notably, however, a variant of the bill color associated SNP chrZ:67480260 (GWAS P = 1.8e^−15^) linked with *TTC39B* does show evidence of selection within subspecies *hecki* (s = 0.0039, logLR = 8.6; Fig 6). This variant is nearly fixed within color-admixed *hecki* (AF = 0.90) and at intermediate frequency within the hybrid zone (AF = 0.41) but – strikingly – is at low frequency within both *acuticauda* (AF = 0.03) and red-billed *hecki* (AF = 0.03). The present geographic distribution of this variant and its inferred genealogy strongly suggests that it did not originate in allopatric populations of either subspecies but instead arose in what are now populations of color-admixed *hecki* (Fig. 6B). The genetic background on which this variant is found is definitively *acuticauda*. Within populations of color-admixed *hecki*, this variant is strongly linked with the *acuticauda* allele of the SNP on the Z chromosome most significantly associated with bill color variation (chrZ:67547840, r^2^ = 0.80, D’ = 0.92) and with the SNP exhibiting greatest genomic cline evidence of introgression from *acuticauda* into *hecki* (chrZ:67498406, r^2^ = 0.87, D’ = 0.96). Allele trajectories indicate that the derived variant at chrZ:67480260 has rapidly increased in frequency between 3 kya and the present day (Fig. 6C). We hypothesize that the SNP showing evidence of a selective sweep from ARG-based inference (chrZ:67480260) arose on – and is currently hitchhiking with – the much older *acuticauda* haplotype of *TTC39B* (represented by chrZ:67498406 and chrZ:67547840) that is the actual target of selection within subspecies *hecki* (see Fig. 6 and Fig. S8).

**Fig. 6.**
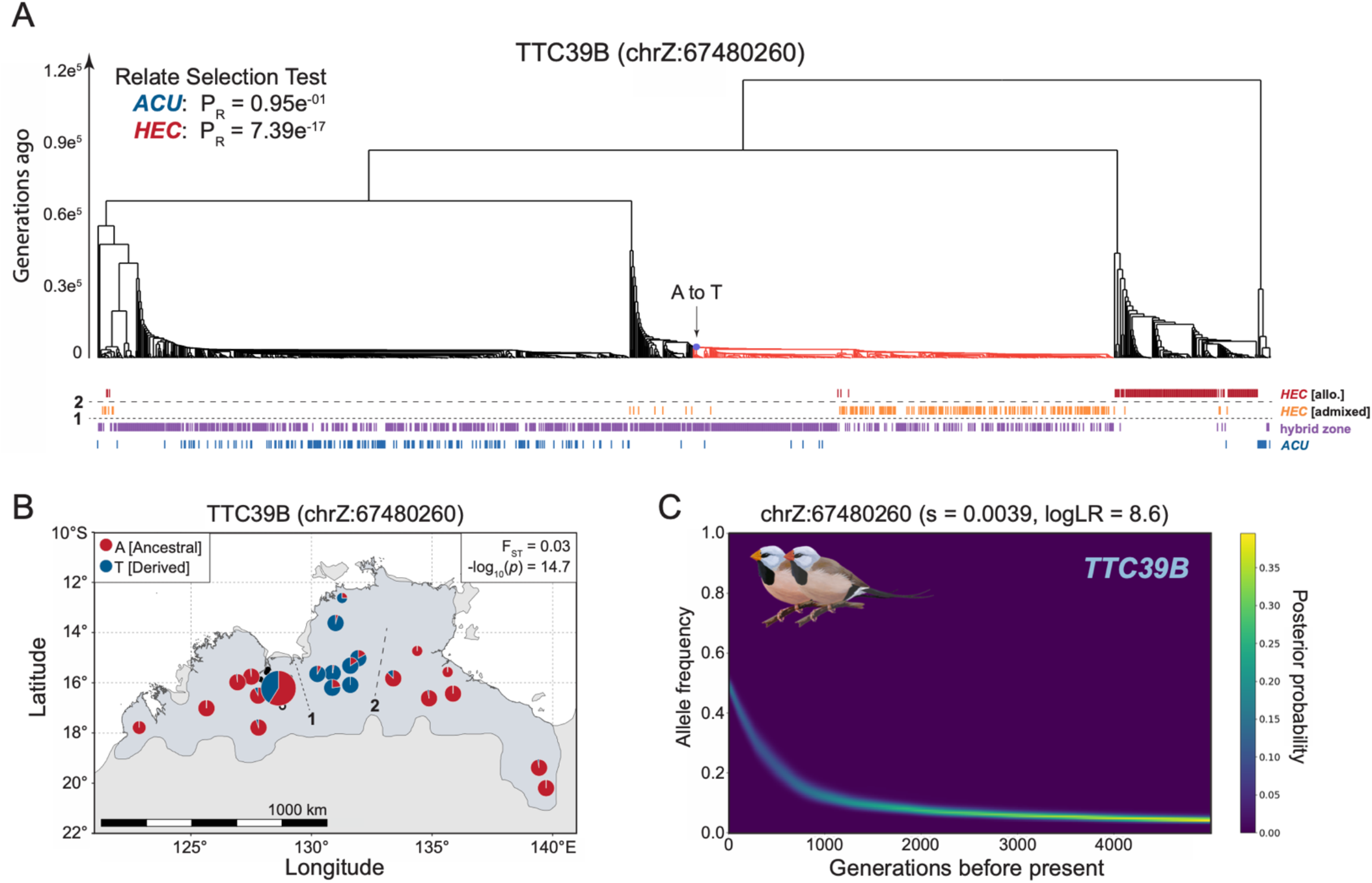
Evidence of selection on *TTC39B* from ancestral recombination graph (ARG) inference. (A) Relate marginal tree for bill color variation associated SNP chrZ:67480260 (2N = 1206 haplotypes). Branches colored in light red represent haplotypes carrying the derived allele at this site. Relate Selection Test evidence for selection over the lifetime of the mutation (P_R_) shown in top right for subspecies *acuticauda* (*ACU*: pops. 1-7, 2N = 178) and subspecies *hecki* (HEC: pops. 20-34, 2N = 312). Vertical hash marks beneath tips of marginal tree represent haplotypes found in males (i.e., with two copies of the Z chromosome) from allopatric red-billed *hecki* (dark red, pops. 28-34), color-admixed hecki (orange, pops. 20-27), the hybrid zone (purple, pops. 8-19), and allopatric yellow-billed *acuticauda* (pops. 1-7). Horizontal dashed lines 1 and 2, as seen in panel (B), represent geographic breaks between the hybrid zone and color-admixed *hecki* (line 1) and between color-admixed and red-billed *hecki* (line 2). (B) Population allele frequencies for SNP chrZ:67480260. Derived and ancestral alleles are color-coded blue and red, respectively. F_ST_ and GWAS significance given in top right inset of panel. (C) Historical allele trajectory of the derived allele at chrZ:67480260 in subspecies *hecki*, which is highly linked with acuticauda-derived SNPs associated with carotenoid ketolation enhancer gene *TTC39B*. A generation time of two years was used to infer timing of selection. s, selection coefficient.

## Discussion

Carotenoid coloration, and particularly coloration produced from metabolized carotenoids, serves as a signal of individual condition in many bird species (Hill and McGraw 2006; Weaver et al., 2018). At the same time, visual signals including carotenoid coloration are hypothesized to serve as important markers of species recognition, competition between males, and mate choice in birds (West-Eberhard 1983; Price 1998; Seddon et al., 2013; Gomes et al., 2016; Price-Waldman et al., 2020) and some experimental work supports these hypotheses (Hill and McGraw 2004). These are patterns observed in contemporary species, but the evolutionary paths to them are only now beginning to be understood. Insights into the evolution of carotenoid ornamentation from natural systems are critical because color variants observed in avian avicultural populations are dominated by highly pleiotropic loss-of-function mutations that are unlikely to persist in the wild (Toews et al., 2017; Price-Waldman et al., 2020; Toomey et al. 2022a).

Here, we report the molecular composition, genetic architecture, and evolutionary history of naturally occurring bill color variation between hybridizing subspecies of the long-tailed finch. We identified C(4)-oxidation as the metabolic process differentiating the two subspecies. The loss of red bill coloration in subspecies *acuticauda* has been achieved via a regulatory change in C(4)-oxidation that halts production of red pigments in the bill but not in the retina. We find that variation in bill hue is most strongly associated with a small number of genes that include *CYP2J19*, an enzyme required for C(4)-oxidation of dietary carotenoids into red ketocarotenoids, and *TTC39B*, a known enhancer of carotenoid metabolism. Genealogical reconstructions indicate that a red-billed phenotype is ancestral in this species. Allelic variation inducing yellow rather than red bills is ancient for both *CYP2J19* and *TTC39B* – on the order of hundreds of thousands of generations – and derived within yellow-billed *acuticauda*. The allelic forms of *CYP2J19* and *TTC39B* that enable yellow bill coloration exhibit signatures of adaptive introgression from *acuticauda* into red-billed *hecki* between 5 kya and the present day. Evidence of selective sweeps on carotenoid processing genes suggest that yellower bill color is favored in the long-tailed finch. Importantly, our evidence argues against a simple alternative frequency-dependent selection (or the related assortative mating) scenario where the most frequent local morph is preferred. Such a scenario is inconsistent with the eastward introgression of the yellow-billed alleles across multiple loci. Taken together, we show that evolutionary transitions between yellow and red color ornamentation, which occur frequently in birds (Ligon et al., 2016; Friedman et al., 2014), can be achieved by natural selection acting upon regulatory mutations of large effect on a small set of genes involved in carotenoid metabolism.

The presence of ketocarotenoids in red cone oil droplets of the retina suggests that subspecies *acuticauda* evolved a yellow bill by suppressing C(4)-oxidation specifically within the bill integument. This simple observation belies a much greater significance for understanding the evolution of carotenoid-based color ornamentation in birds more broadly. Nearly all diurnal birds have red cone oil droplets in their retinas, which means that they can synthesize red ketocarotenoids from dietary carotenoid precursors, but comparatively few have red bills or feathers (Goldsmith et al., 1984; Vorobyev 2003). Indeed, it has been posited that carotenoid ketolation in reptiles originally evolved in the context of color vision before only later being co-opted to produce color ornamentation (Twyman et al., 2016). In groups of birds with carotenoid-based plumage and bill coloration, transitions between yellow ornamentation produced by modified yellow carotenoids and red ornamentation produced from modified red carotenoids can often evolve dynamically (Ligon et al., 2016; Friedman et al., 2014) but the underlying genetic mechanisms responsible have rarely been characterized (Twyman et al., 2018). One working hypothesis is that such evolutionary transitions can be achieved by regulatory changes to where and when *CYP2J19* expression occurs [while leaving *BDH1L* expression unchanged] (Toomey et al. 2022a). Consistent with this regulatory hypothesis, we observed an abundance of strongly associated upstream variants and a lack of coding level differences between *acuticauda* and *hecki* alleles of *CYP2J19*. We found no evidence that *BDH1L* was associated with color variation in this system. We posit that the evolutionary transition from red to yellow bill coloration in the long-tailed finch has arisen largely through regulatory changes to where expression of *CYP2J19* – and thus C(4)-oxidation – occurs. This mechanism may be widely responsible for the frequent evolutionary transitions between carotenoid-color ornamentation observed in closely related species.

Identifying which tissues are involved in carotenoid ketolation is a critical step for evaluating any potential benefits or costs associated with the production of ketocarotenoid ornamentation. In avian species with red feathers, *CYP2J19* activity appears to predominately occur in the liver while activity is largely restricted to the periphery in species with red bare parts (i.e., bill and legs) (Alonso-Alvarez et al., 2022). However, variation in the location of carotenoid metabolism exists even between species with similar ornamentation. For example, both the red-billed quelea *Quelea quelea* and the zebra finch have ketocarotenoid-rich red bills and tarsi but have distinct modes of producing them: in the former, carotenoid metabolism is concentrated in the liver and ketocarotenoids are then shuttled to their site of deposition while, in the latter, ketolation is absent from the liver and instead appears restricted to the peripheral sites of carotenoid deposition (Twyman et al., 2018; Mundy et al., 2016). In the zebra finch, this regionalization of carotenoid metabolism is achieved in part via a tandem duplication of *CYP2J19*; with one copy expressed only in the retina and the other in both bill and retina (Mundy et al., 2016). We find no evidence for a second copy of *CYP2J19* in the long-tailed finch (see Materials and Methods) and only one copy has been found in other species of bird examined to date (Twyman et al., 2018; Emerling et al., 2018). Further work is therefore required to ascertain the biochemical mechanisms by which the long-tailed finch (namely subspecies *acuticauda*) regulates the metabolism and deposition of carotenoids between different bodily regions.

Both *CYP2J19* and *TTC39B*, the two genes that together explain 40.5% of variance in bill hue, exhibited clear signatures of adaptive introgression from yellow-billed *acuticauda* into red-billed *hecki*. Recent experimental work shows that *CYP2J19* and *TTC39B* interact epistatically, potentially by *TTC39B* modulating the rate of transport of carotenoid substrates to or from the site of enzymatic conversion by *CYP2J19* (Toomey et al., 2022a). An epistatic relationship between these two genes appears evident in the long-tailed finch (Fig. 4F). For individuals from an otherwise *hecki* genomic background (i.e., from pops. 20 – 34, N = 234), a linear model containing genotype information for both loci provided a significantly better fit for bill hue variation than models containing only one locus (ΔAICc > 45). We therefore predict that the yellow bills of subspecies *acuticauda* are largely a result of an absence of *CYP2J19* expression and a suppression of *TTC39B* expression within the bill integument (Fig. 7A).

**Fig. 7.**
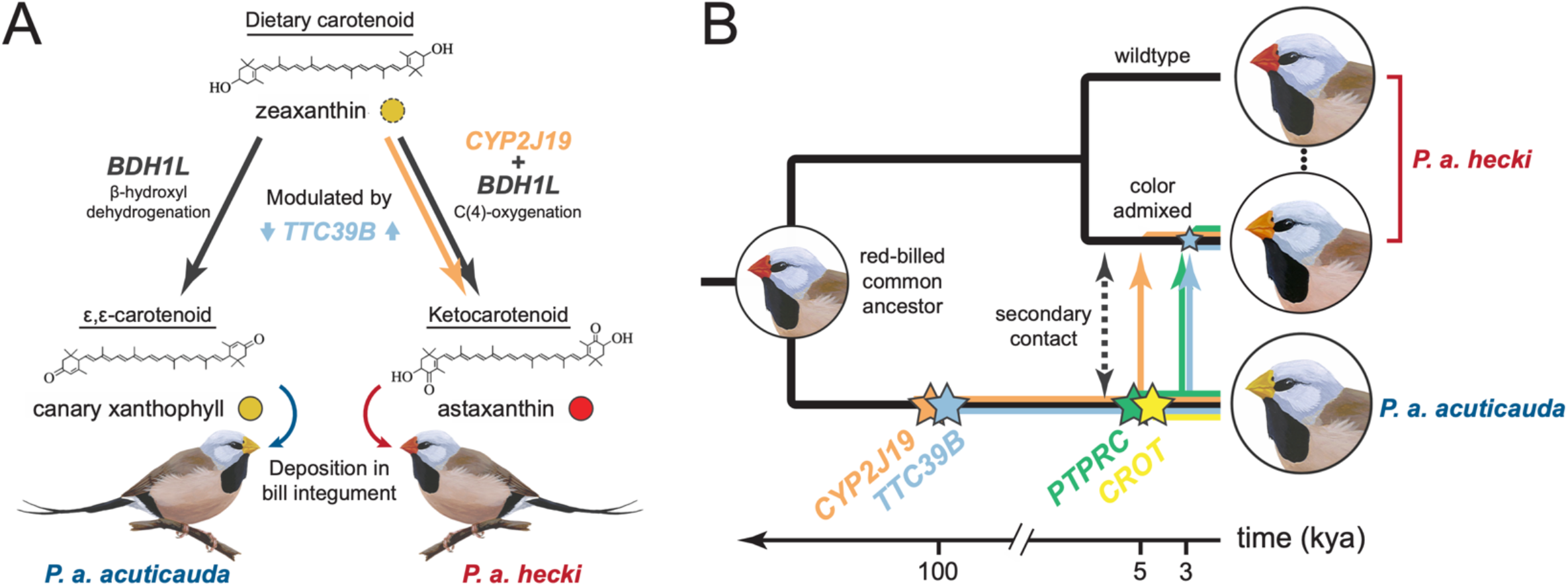
Summary of bill color regulation and evolution in the long-tailed finch. (A) In birds and fish, the production of red ketocarotenoids requires the enzymatic activity of both *CYP2J19* and *BDH1L* while the production of ε,ε-carotenoids occurs without *CYP2J19* (Toomey et al. 2022a). The absence of ketocarotenoids in the bill integument of yellow-billed *acuticauda* is most likely due to a lack of expression of *CYP2J19*, which thereby precludes C(4)-oxidation of dietary carotenoids like zeaxanthin into red ketocarotenoids like astaxanthin. The detection of canary xanthophylls, a class of ε,ε-carotenoid, in yellow bills suggest that *BDH1L* – and not *CYP2J19* – is enzymatically active in *acuticauda*. We predict that the *acuticauda* allele of *TTC39B*, a known modulator of carotenoid metabolic efficiency (Toomey et al. 2022a), suppresses the rate of carotenoid metabolism while the *hecki* allele enhances it. The contribution to carotenoid processing of *PTPRC* and *CROT* are not yet known. (B) Hypothesized evolutionary history of four loci in genomic regions associated with bill color variation exhibiting evidence of selective sweeps. Colored stars represent the approximate age and lineage of origin of each allele as inferred by ARG-based inference: orange (*CYP2J19*), blue (*TTC39B*), green (*PTPRC*), and yellow (*CROT*). The polarization of derived change within yellow-billed *acuticauda* suggest that the most recent common ancestor of long-tailed finch subspecies had a red bill. The alleles most strongly associated with bill color variation at *CYP2J19* (e.g., chr8:3144828) and *TTC39B* (e.g., chrZ:67547840) arose approximately 100 kya in subspecies *acuticauda*, presumably during a period of allopatric divergence. Alleles at both loci have subsequently introgressed into subspecies *hecki* following secondary contact, and they have rapidly increased in frequency between 5 and 3 kya and the present day, respectively. The small blue star on the branch representing color-admixed *hecki* denotes the *TTC39B* variant at chrZ:67480260 that bears the signatures of a selective sweep on an *acuticauda* haplotype background. Alleles associated with *PTPRC* first underwent a rapid increase in frequency within *acuticauda* approximately 5 kya before introgressing into subspecies *hecki* and rapidly increasing in frequency between 3 kya and the present day. Selective sweeps on *acuticauda*-derived alleles at *TTC39B* and *PTPRC* appear to have initiated in *hecki* at approximately the same time (~3 kya). Evidence suggests that a selective sweep in *acuticauda* on alleles within the gene *CROT* occurred between 4 kya and the present day and have not introgressed across the long-tailed finch hybrid zone.

Clinal evidence of introgression of yellow alleles derived in *acuticauda* into an otherwise *hecki* genomic background is clear. What evolutionary forces propelled their spread? Selective sweep analyses confirm a history of natural selection on bill color in the long-tailed finch. All four of the top association peak regions carry the hallmark signatures of selective sweeps. Allelic variation derived within yellow-billed *acuticauda* and associated with *CYP2J19*, *TTC39B*, and *PTPRC* have increased in frequency in *hecki* under moderate to strong selection (i.e., s > 0.003) between 5 kya and the present. Evolutionary reconstructions suggest that allelic divergence between subspecies at both *CYP2J19* and *TTC39B* is ancient: gene trees including the two SNPs most strongly associated with bill color variation coalesce approximately 100 kya (Fig. 7B; Fig. S5, Fig. S8). The temporal discrepancy between the age of *CYP2J19* and *TTC39B* alleles (on the order hundreds of thousands of generations) and the inferred timing of selection on them within *hecki* (on the order of thousands of generations ago) is consistent with a model of allopatric divergence between long-tailed finch subspecies in Pleistocene refugia that preceeded introgression following secondary contact (Bowman et al. 2010; Fig. 7B). Evidence of natural selection associated with *PTPRC* was initiated much more recently: first within subspecies *acuticauda* 4-5 kya and then within color-admixed populations of *hecki* between 3 kya and the present (Fig. 7B; Fig. S6, Fig. S10). In contrast, the signature of a selective sweep centered on the gene *CROT* has been restricted to subspecies *acuticauda* and occurred between 4 kya and the present (Fig. S7, S11, and S15). Consistent with lines of evidence from geographic and genomic cline analyses, the rapid increase in frequency of allelic variation associated with bill hue suggests that the spread of yellow coloration has been adaptive.

One tantalizing hypothesis to explain the spread of yellow bill color alleles is that long-tailed finches – of both subspecies – prefer partners with yellower bills. This argument for a role of sexual selection, however, has little empirical support. A behavioral assay performed in controlled captive conditions found that female long-tailed finches exhibit an assortative mate preference: i.e., red-billed females prefer to spend time with red-billed males over yellow-billed males, and vice versa (McDiarmid et al., 2023). Moreover, experimental manipulation of bill color, done to control for potential differences between subspecies in song or other aspects of male courtship behavior, eroded the strength of assortative mate preference to non-significance. Together this suggests that bill coloration is just one trait of many used by these birds to evaluate and select a mate. Long-tailed finches used in these experiments were sourced from allopatric populations of each subspecies where bill color variation is low (see Fig. 1D) and so they may potentially not have been well-suited to address whether bill color affects mate choice or mating success. It would be more appropriate to address these questions in nature using populations with the greatest individual variation in bill color (e.g., color-admixed pops. 20 – 27).

Color ornamentation differences – and sometimes little else – can be key components of reproductive isolation between closely related bird species (Toews et al., 2016; Campagna et al., 2017; Stryjewski and Sorenson 2017). In the long-tailed finch, bill coloration does not appear to play such a role. Rather than acting as a trait that inhibits genetic exchange, introgression of yellow bill color alleles appears to have been favored by natural selection and has progressed to such an extent that the genomic hybrid zone between long-tailed finch subspecies is phenotypically cryptic. Indeed, the recent increase in frequency of yellow bill color alleles within *hecki* may plausibly be replicating today the very selective sweeps that drove these alleles to fixation within *acuticauda* long ago. Evidence of introgression of genes associated with ornamentation differences between allopatric populations is not without precedent and many cases have been associated with sexual selection. In birds, the asymmetrical introgression of color alleles between hybridizing taxa due to mate preference for the introgressing phenotype has been reported in fairywrens (Baldassarre et al., 2014), mannikins (Parsons et al., 1993; McDonald et al., 2001; Stein and Uy 2006), tinkerbirds (Kirschel et al., 2020), and wagtails (Semenov et al., 2021). A particularly striking example of this in deeper time comes from new world warblers in the genus *Setophaga*, where alleles of the carotenoid-cleaving beta-carotene oxygenase 2 gene (*BCO2*) conferring yellow plumage show evidence of having repeatedly been exchanged between distantly related species (Baiz et al., 2021). Determining the fitness effects of divergent metabolic programs for the processing of carotenoids, which ultimately manifests as yellow or red bill coloration in long-tailed finches, will be key to understanding the putatively adaptive evolution of bill coloration in these birds. The increasing accessibility of linked-read population-scale genomic data provides evolutionary biologists with a better toolkit than ever before with which to examine the genes, evolutionary history, and role of diversifying selection underlying the diversity of colorful ornamentation of birds and other animals.

## Materials and Methods

### Animal care and use

All experimental procedures described in this study were approved by the applicable ethics committee or authorities at Macquarie University, Australia. The study and sampling of finches has been approved by the Australian government. Long-tailed finch *Poephila acuticauda* individuals of each subspecies were bred in captivity from wild-derived stocks established less than ten generations prior. The population of subspecies *acuticauda* was descended from individuals collected from two locations in Western Australia in 2009: Mt. House (17°02’S, 125°35’E) and Nelson’s Hole (15°49’S, 127°30’E). The population of subspecies hecki was descended from individuals collected from October Creek (16°37’S, 134°51’E) in the Northern Territory in 2010. Populations of each subspecies were kept reproductively isolated from one another save for the purpose of experimentally producing F_1_ hybrids.

### Bill color phenotyping

Bill color was measured via UV-vis reflectance spectrophotometry for wild-caught and captive-bred adult long-tailed finches following protocols published in prior studies (Griffith and Hooper 2017; McDiarmid et al., 2023). In brief, spectral reflectance of the upper mandible from three consecutive scans per individual were averaged and smoothed using the R package Pavo 2 (Maia et al., 2019). Spectra were then normalized by their maximum and minimum reflectance values, and the colorimetric variable H3 was calculated for all samples. H3 is a measure of bill hue that represents the wavelength midway between the minimum and maximum reflectance of a surface, which we bounded between 400 and 700 nm (Maia et al., 2019) and has previously been shown to effectively differentiate the bill colors of the two long-tailed finch subspecies and their hybrids (McDiarmid et al., 2023). To supplement reflectance data first examined by Griffith and Hooper (2017) and McDiarmid et al. (2023) we measured bill color for an additional 165 wild-caught and 137 captive-bred samples in this study for a total of 948 wild and 550 captive samples with reflectance data (N = 1498 total).

### Evaluation of bill carotenoid composition with HPLC

Carotenoids were isolated from the bill tissue of five individuals of each subspecies following procedures adapted from McGraw et al. 2002 and McGraw and Toomey, 2010. Thin slices of integument (0.002-0.01 g) were shaved from the outer bill using a razor. Carotenoids were extracted from the shavings in the presence of solvent (5–6 mL hexane: tert butyl methyl ether, 1:1, v/v) using a mortar and pestle. Ground tissue and solvent were centrifuged, and the supernatant recovered for saponification to remove esterification that impedes elution and accurate quantification of carotenoids via HPLC (McGraw and Toomey, 2010). Importantly, while this method increases recovery of ketocarotenoids and dietary carotenoids, it does result in the loss of canary xanthophylls a and b (Toomey et al. 2022b). To saponify carotenoid samples, the supernatant was evaporated to dryness and the carotenoids resuspended in 100 µL of 100% ETOH. Next, we added 100 µL of 0.02M KOH in MeOH, vortexed for 30 seconds, capped with N_2_ gas, and incubated the extract at RT in the dark overnight. After incubation, 250 µL H_2_0, 500 µL TBME (100%), and 250 µL hexane (100%) were added sequentially, with vortexing after the addition of the H_2_0 and hexane. Finally, esters were precipitated from the extract with the addition of 100 µL saturated saltwater (pure NaCl) and 30 seconds of vortexing. The saponified carotenoids were moved to a new tube and evaporated to dryness before resuspension in 200 µL of acetone for immediate HPLC injection.

Carotenoid samples were analyzed via HPLC using a Shimadzu Prominence UPLC system. Extracts were injected in 10 µL volumes into a Sonoma C18 column (10 µm, 250 x 4.6 mm, ES Technologies, New Jersey, USA) fitted with a C18 guard cartridge. Separation and elution of carotenoids was done using an adapted tertiary mobile phase (adapted from Wright et al. 1991). The mobile phases used here were: A) 80:20 methanol: 0.5M ammonium acetate; B) 90:10 acetonitrile: H2O; and C) ethyl acetate. We used a tertiary linear gradient with a flow rate of 1mL min^−1^ that consisted of 100% A to 100% B over 4 min, then 80% C: 20% B over 14 minutes, then 100% B over 3 minutes, ending with 100% A over 11 minutes to re-equilibrate the column. Samples were run consecutively with an autosampler fitted with an internal cleaning port with 1:1 v/v MeOH:H_2_O to remove cross-contamination. Carotenoids were detected using a Prominence UV/Vis detector set to 450 nm. Carotenoids were identified based on comparisons to pure standards, an internal system database of retention times, and published accounts.

### Retinal imaging and cone photoreceptor subtype classification

Eyes were obtained from frozen and/or freshly deceased long tailed finches that had been culled for a complementary project. Three yellow-billed *P. a. acuticauda* and three red-billed *P. a. hecki* birds were examined. The left eye of each finch was removed and hemisected at the equator using a scalpel blade. The posterior segment of the eye was placed into phosphate buffered saline (PBS; pH 7.4; 340 mOsmol kg^−1^) to facilitate dissection of the retina. Small pieces (~2×2 mm) of retina from the dorsal or ventral retinal periphery were dissected away and mounted on a glass microscope slide in a drop of PBS, and a top coverslip applied and sealed with clear nail varnish. The retina was viewed under bright-field illumination using an Olympus ×100/NA 1.4 oil immersion objective on an Olympus BX-53 microscope fitted with DIC optics. Images were captured using an Olympus DP74 digital camera and cellSens software.

### Reference genome assemblies

Zebra finch *Taeniopygia guttata* v1.4 (GCF_003957565.2).

### Enrichment for high-molecular weight DNA

We performed high-molecular weight (HMW) DNA enrichment to compensate for the high level of degradation in most of the gDNA samples. Here, we chose a cut-off of >8 kbp. Briefly, 50 µl of gDNA in modified Ampure bead buffer (10mM Tris pH=8, 0.1mM EDTA, 18% PEG8000, 2.5M NaCl) was gently mixed with 64 µl of the size selection beads (Cytiva, 65152105050250) in size selection buffer (20 mM Tris pH=8, 6% PEG8000, 833 mM NaCl, 70 mM MgCl2). Size selection reaction was then incubated 25 min at 65°C. Beads were then washed on magnetic stand twice with 80% ethanol and the enriched HMW gDNA was eluted with TE buffer at 45°C for 30 minutes.

### Sequencing library construction

Linked-read (LR) genomic libraries were prepared for 1229 samples following published protocols from Meier et al. (2021) with the following modifications to the 96-plex “haplotagging” protocol. Briefly, 0.375 ng of each HMW DNA sample was tagged with 1.25 µl of haplotagging beads from an expanded panel of 3 separate 96-well Haplotag bead plates, each carrying a different i7/i5-barcode ligation overhang: “T/G” or “C/T”, in addition to the original “A/C” overhang in (Meier et al., 2021). This expanded panel enabled sequencing of 288 samples (3 x 96) per NovaSeq sequencing lane. Subsampling of beads after tagmentation was carried out at a 1:5.6 ratio to maintain high reads-per-molecules ratio across the individuals. This corresponds to approximately 1.6e^6^ beads (barcodes) to 67 pg DNA per sample, or 135 haploid copies of the long-tailed finch genome. Pooled and subsampled beads of each of the plates, carrying a total of 6.4 ng DNA per library (96 x 67 pg), were split into two equal samples and incubated at 50°C for 25 minutes with exonuclease 1 (NEB, M0293) to remove unintegrated barcoded transposon adapters. Each plate’s DNA library was then amplified in two 50 µl Q5 High-Fidelity DNA Polymerase reactions (New England BioLabs) for 11 thermocycles, then twice size selected with Ampure beads at 0.45× followed by 0.85× bead:sample ratios to remove >1 kbp library and <300 bp fragments, respectively. The set of samples with LR data included 1133 long-tailed finches: 982 from across the geographic distribution of the species in the wild and 171 captive reared individuals descended from wild-caught individuals of each subspecies held in aviaries at Macquarie University in Sydney, Australia. We also prepared LR libraries for 96 black-throated finches *Poephila cincta* (48 *P. c. cincta* and 48 *P. c. atropygialis*). This allopatric sister species to the long-tailed finch was used as a closely related outgroup in population genetic analyses. Bill color phenotype data was available for 508 of the sequenced samples.

Short-read (SR) genomic libraries were prepared for 22 samples (5 *P. a. acuticauda*, 15 *P. a. hecki*, 1 *P. c. cincta*, and 1 *P. c. atropygialis*) using an Illumina TruSeq DNA Library Preparation Kit (Illumina, San Diego). The 20 long-tailed finch samples with SR libraries were used as technical replicates for evaluating genotype imputation accuracy and phasing performance with LR data.

### Sequencing and demultiplexing

LR libraries were sequenced on a NovaSeq 6000 2x150bp S4 flow cell (Illumina, San Diego) from a commercial service provider (MedGenome, Foster City, USA) with a 151+13+13+151 cycle run setting for a total of 3.3 TBases of sequencing data. Raw CBCL sequencing data was converted to fastq using bcl2fastq (Illumina) without sample sheet and with parameters --use-bases-mask=Y151,I13,I13,Y151 --minimum-trimmed-read-length=1 --masked-short-adapter-reads=1 --create-fastq-for-index-reads. beadTag demultiplexing was performed using a demultiplexing c++ script to decode each of the four barcodes and saved as BX:Z tag information in modified fastq files. Reads were subsequently split into plates based on the combination of ligation overhangs. Demultiplexing information is available at https://github.com/evolgenomics/haplotagging. Reads were trimmed to remove adapter sequences and low-quality bases (Cutadapt; https://doi.org/10.14806/ej.17.1.200), mapped to the zebra finch reference genome using BWA mem (Li 2013), and PCR duplicates removed under barcode-aware mode Picard’s MarkDuplicates module (http://broadinstitute.github.io/picard/). We recovered an average of 8.4 million paired end reads and a median depth of coverage of 1.38× for the 1229 samples with LR data.

We included data from 20 additional wild-caught long-tailed finch samples generated in a previous study (10 *P. a. acuticauda* and 10 *P. a. hecki*; standard SR genomic sequence from ENA Study: PRJEB10586; Singhal et al., 2015). Reads (2x150) from all SR data samples were processed as above, except without barcode-aware duplicate marking. The 42 samples with SR sequence data had an average of 150 million paired end reads and a mean depth of coverage of 26.1×.

### Variant discovery and genotyping

We generated an initial set of variants with bcftools mpileup using the full set of 1249 samples. A subset of biallelic SNVs were then selected after bcftools filtering for proximity to indels (i.e., −g3) and based on variant quality and sequencing error artefacts (%QUAL<500 || AC<2 %QUAL<50 || %MAX(AD)/%MAX(DP)<=0.3 || RPB<0.1 && %QUAL<50). This quality-filtered subset of variants was then pruned of sites that overlapped with annotated repetitive regions of the zebra finch genome called by repeatMasker (Smit et al. 2013-2015; Flynn et al. 2020). A total of 37.69 million SNVs remained after initial quality filtering and repeat masking.

Genotype calling and imputation with this set of SNVs was performed using read data in BAM format from all 1249 samples in 1 Mb windows using STITCH version 1.6.6 (Davies et al. 2016) in pseudohaploid mode with the following parameters: K=100, nGen=1000, shuffle_bin_radius=100, niterations=40, switchModelIterations=25, buffer=50000. Based on the significant negative relationship between chromosome size and per bp recombination rate in birds (Singhal, Leffler et al. 2015), STITCH was run across three chromosome classes using the following recombination rate tuning (i.e., >25 Mb, expRate = 1.0; <25 Mb, expRate = 5.0, and < 10 Mb, expRate = 10.0). Microchromosomes smaller than 2.6 Mb (a set of 9 chromosomes encompassing <1.5% of the genome) were excluded from further analysis due to poor imputation performance likely resulting from their high per bp rate of recombination. A final set of 33.23 million SNVs with high information content (INFO_SCORE > 0.4) were retained for downstream analyses (88.2% of the initial set of variants identified).

### Population genetic analyses

We calculated F_ST_ between allopatric populations of each subspecies (i.e., 110 *acuticauda* from pops. 1 – 7 and 110 *hecki* from pops. 28 – 34 in Figure 1) for all SNPs with a minor allele count of at least 2 and in 20 kb windows along each chromosome with a step size of 10 kb using VCFtools version 0.1.16. We quantified F_ST_ on the Z chromosome using a subset of allopatric males (70 *acuticauda* and 69 *hecki* samples) to circumvent any effects of female hemizygosity. We investigated fine-scale patterns of introgression between subspecies by calculating ΔF_ST_ in 10 kb windows with 5 kb steps using a set of color admixed samples from pops. 20 – 27 as follows:

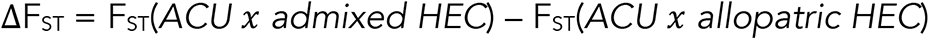

After removing windows with less than 50 variants on each chromosome (and less than 20 variants on the Z chromosome) we classified genomic regions in the 0.5^th^ and 99.5^th^ percentile of the ΔF_ST_ distribution as showing potential evidence of introgression from *acuticauda* into *hecki* and from *hecki* into *acuticauda*, respectively.

We performed hybrid index estimation using a set of linkage-disequilibrium (LD) pruned ancestry informative SNP markers. We defined a set of ancestry informative markers as the 755 autosomal SNPs above an F_ST_ threshold of 0.7 and the 42668 Z chromosome SNPs above an F_ST_ threshold of 0.95 between allopatric populations. We used PLINK v.1.9 to LD-prune SNPs in this marker set above an r^2^ of 0.2 within 100 kb windows and a 10 kb step size (--indep-pairwise 100 10 0.2) and with any amount of missing data across samples (--geno 0.0). Of the 43423 ancestry informative SNP markers, 1137 SNPs remained after LD pruning. We used the R package gghybrid (Bailey 2023; https://doi.org/10.5281/zenodo.3676498) to perform Bayesian MCMC hybrid index estimation on a set of 649 ancestry informative SNPs remaining after further filtering for an AF difference of at least 0.8 between allopatric populations and a MAF of less than 0.05 in one of the two allopatric populations. This set of markers included 99 autosomal SNPs and 550 Z-linked SNPs. We modeled the ancestry proportion of all 1153 wild and captive long-tailed finch samples as the h posterior mode from gghybrid function hindlabel.

### GWAS

We performed a genome-wide association study (GWAS) for bill color using a measure of bill hue: H3. As described above, the colorimetric variable H3 was quantified from reflectance spectrophotometer readings as the wavelength midway between the minimum and maximum reflectance of light reflected off a bird’s upper mandible. We retained a total of 508 individuals with bill color phenotype data for association mapping. We investigated the genetic basis of bill color using the Wald test implemented in Gemma version 0.98.5 (Zhou and Stephens 2014). We fit a univariate linear mixed model to test for an association between phenotype and each SNP genotype. We included a relatedness matrix to correct for population structure and because hybrid sex (McDiarmid et al. 2023) and genome-wide hybrid index are significantly correlated with bill hue (Spearman’s r = 0.66, p < 2.2×10^−16^) we also included these as covariates during modeling. Keeping only variants with a minor allele frequency above 2%, genome-wide data from a total of 17,566,649 variants were analyzed (mean of 17.3 SNPs / kbp).

We evaluated the genome-wide threshold for significance at a false discovery rate of 1% (α = 0.01) in two ways. First, using a Bonferroni correction on the total number of SNPs tested (threshold = 9.24; P = 5.69e^−10^) and second, as the 99^th^ percentile across the distribution of differences following 1,000 permutations of bill hue onto observed genotypes (threshold = 9.03; P = 9.29e^−10^). As results were qualitatively identical with respect to using either significance threshold approach (523 versus 576 significant SNPs), we adopted the permutation derived threshold as it preserves important features of the data while making fewer assumptions. Results were visualized by log-transforming the p-values, changing their signs, and generating Manhattan plots using the R package fastman (https://github.com/kaustubhad/fastman).

We defined association peaks as clusters of at least five SNPs above our genome-wide significance threshold that were separated by <50 kb from the next significant SNP. We scanned for protein-coding genes within 100 kb upstream and downstream of each association peak in R using the annotation file for the zebra finch reference genome (GCF_003957565.2). Of the eleven identified association peaks (Table 1), we focused subsequent attention on the three autosomal peaks and the single Z chromosome peak that contained the most SNPs significantly associated with color variation genome-wide, included SNPs observed to be highly differentiated between subspecies, and encompassed genes previously associated with or plausibly linked to carotenoid-based color variation. The functional significance of all SNPs above genome-wide significance was evaluated using SnpEff v5.2a (Cingolani et al. 2012).

Following Zhou et al. (2013), the proportion of genetic variance explained (i.e., PVE) by the most strongly associated SNP variant in each association peak was estimated as:

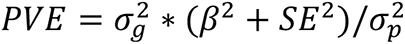

Where genotypic variance was represented by 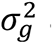 and phenotypic variance by 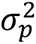, respectively. Estimates of allelic effect size (*β*) and its standard error (*SE*) were inferred for each SNP by GEMMA.

### CYP2J19 copy number evaluation

The study of the zebra finch that first identified *CYP2J19* as a candidate oxidative ketolation enzyme (Mundy et al., 2016) reported two copies of this gene on chromosome 8. The authors proposed that the second copy had arisen due to tandem duplication with one specialized for color vision (denoted *CYP2J19A*) and the other for color ornamentation (*CYP2J19B*) (Mundy et al., 2016). To our knowledge, only a single copy of *CYP2J19* has been detected in any other avian genome (Twyman et al. 2018; Emerling 2018) and only a single copy of CYP2J19 is present in the zebra finch reference genome (GCF_003957565.2). We examined whether there is any evidence of two copies of *CYP2J19* in the long-tailed finch and black-throated finch by evaluating deviations in depth of coverage along chromosome 8. Specifically, we evaluated whether samples exhibited an increase in depth of coverage in 1 kb sliding windows across *CY2PJ19* (chr8:3167181-3178161) that would indicate that reads from two copies of this gene were mapping to the same genomic location: a ~2× increase in coverage would be expected if this was the case. We did not detect any increase in depth of coverage for reads mapping to *CYP2J19* in the zebra finch reference genome that might indicate this gene exists in more than one copy in the genus *Poephila*.

### Geographic clinal analysis

One-dimensional Bayesian analyses of cline position and shape were performed using a subset of 30 transect populations with at least 2N = 10 samples available using the R package BAHZ (Thurman 2019; https://github.com/tjthurman/BAHZ). Geographic clines were modeled for the most strongly associated SNP in each of the four focal GWAS association peaks and for genome-wide hybrid index. We determined the best-fitting cline model based on changes in allele frequencies across our sampling transect using two parameters, cline center and width.

### Genomic clinal analysis

We evaluated our set of 649 ancestry informative markers in linkage equilibrium (see population genetic analyses above) for evidence of biased introgression between long-tailed finch subspecies using the ggcline function of gghybrid (Bailey 2023). We used our genome-wide hybrid index and genotype data from 982 wild-caught samples to model the genomic cline center (*c*) and steepness (*v*) for each SNP marker.

### Phasing

Molecular phasing of SNPs on chromosomes 2, 8, and Z was performed using haplotagging LR molecular barcode information with HapCUT2 (Edge et al. 2017). Phasing was performed using a BAM file containing LRs mapped to the zebra finch reference genome and a VCF file containing variant calls and diploid genotypes for the same individual. As female samples are hemizygous for the Z chromosome, and as a result are expected to already be phased apart from the pseudo-autosomal region, we did not include them in our HapCUT2 pipeline. Phasing performance, measured as N50 haplotype length, was evaluated as follows. We first compared the distribution of individual N50 haplotype lengths for each chromosome between the 1229 samples with LR data and low sequencing depth (e.g., 1.4×) against the 40 samples with standard short-read data and deep sequencing depth (e.g., 30.0×). We next directly compared N50 haplotype lengths for the 20 individuals in our dataset that were sequenced using both linked-read and short-read approaches to control for the effects of sample quality.

### Selection

Scans for signatures of selection were performed on four regions of the genome containing the GWAS association peaks most strongly associated with variation in bill color. We defined each region as a genomic window 1 Mb downstream and upstream of each association peak. The mean size of each association peak was 0.13 Mb (range from 0.03 to 0.33 Mb) while the mean genomic window evaluated was 2.13 Mb (range from 2.03 to 2.33 Mb), respectively. This approach allowed us to evaluate evidence of selective sweeps at each association peak against the genomic background they were located upon. Evidence of positive selection was evaluated based on summary statistics of haplotype structure and Ancestral Recombination Graph (ARG) inference to test for and estimate the strength and timing of selection, as well as estimate the full allele frequency trajectory.

We evaluated long-range haplotype homozygosity within populations using the integrated haplotype homozygosity score (iHS) and compared haplotype homozygosity between populations using cross-population extended haplotype homozygosity (xpEHH). Both statistics were calculated with phased genotype data using the R package rehh (Gautier et al., 2017; Klassmann and Gautier 2022). We focused on three groups of sampled populations for calculating summary statistics based on haplotype structure: (*i*) allopatric yellow-billed *acuticauda* (pops. 1 – 7), (*ii*) color-admixed *hecki* (pops. 20 – 27), and (*iii*) allopatric red-billed *hecki* (pops. 28 – 34). We did not include hybrid zone populations (pops. 8 – 19) in these calculations because of their demographic composition of recent generation hybrids.

We leveraged the LR supported phasing in our dataset using RELATE version 1.1 (Speidel et al., 2019) to examine evidence of positive selection at the four genomic regions encompassing bill color association peaks. We first converted phased VCFs to the haps/sample file format used by Relate with the RelateFileFormats.sh script and prepared the input files using PrepareInputFiles.sh (both part of the Relate package). We ran Relate on 1928 haplotypes from males and females for autosomal chromosomes 2 and 8 and on 1206 haplotypes from males for the Z chromosome. We used the zebra finch reference sequence (GCF_003957565.2) to polarize variants as ancestral or derived. We ran Relate with options -m 6.25e-9 -N 5e5 for the autosomal chromosomes and -m 6.94e-9 -N 2e5 for the Z chromosome to account for the difference in germline mutation rate and effective population size between these chromosome classes (de Manuel et al., 2022; Bergeron et al., 2023). We supplied genetic maps for each chromosome generated from genotype calls from 70 allopatric *acuticauda* males using ReLERNN version 1.0.0 (Adrion et al., 2020), a deep learning approach that uses recurrent neural networks. ReLERNN was run using the simulate, train, predict, and bscorrect modules with default settings apart from the mutation rates specified above and a generation time of two years. Inferred recombination rates were averaged in 1 Mb blocks in 50 kb sliding windows. We used the Relate script EstimatePopulationSize.sh with options -m 6.25e-9 (autosomal) or -m 6.94e-9 (chrZ) and -years_per_gen 2.

We extracted the genealogies of each of our focal groups (i.e., *acuticauda* pops. 1-7 and *hecki* pops. 20-34), re-estimated population size history for them, and used the output of the previous step as input for the Relate Selection Tests script DetectSelection.sh. This approach tests for evidence of positive selection on a particular variant based on the speed at which lineages carrying it spread relative to other ‘competing’ lineages (Speidel et al., 2019). We accounted for the demographic history of each group using the same tuning for mutation rate and generation time differences between chromosome classes as above. From the .sele files generated, we extracted p-values from the column “when_mutation_has_freq2” which tests for evidence of selection over the lifetime of a particular variant.

We further investigated the evolutionary history of each variant found to show evidence of positive selection within our four association peak regions using CLUES (Stern et al., 2019). We used the Relate script SampleBranchLengths.sh on the genealogies of each of our groups to sample branch lengths from the posterior and account for uncertainty. We ran 100 samples (-num_samples 100) and accounted for the demographic history of each group using the .coal files output from EstimatePopulationSize.sh. We then ran CLUES (inference.py script) with the option -coal to account for demographic history. We fine-mapped variants using the likelihood ratio statistic generated by CLUES as suggested by Stern et al. (2019) and focused on the variants within each association peak that show moderate to strong selection (s > 0.003). We used the plot_traj.py script to plot the results. As evidence from population genetic, geographic cline, and genomic cline analyses together suggested that allelic variation associated with bill color variation has predominately introgressed from subspecies *acuticauda* into subspecies *hecki*, we focused attention on estimating the strength and timing of selection, as well as the full allele frequency trajectory, within subspecies *hecki* (i.e., pops. 20-34).

## Data and Code Availability

All analytical code will be deposited in GitHub, https://github.com/dhooper1/Long-tailed-Finch. The genomic data will be archived in GenBank (BioProject ID PRJNAXXXXXX). Phenotypic and collection data for all *Poephila* samples are given in Supplemental Tables.

## Acknowledgements

This work was funded by ARC-DP-180101783 to DMH and SCG and NSF-IOS-2037741 to GEH. DMH was supported by the Gerstner Scholars Fellowship and the Gerstner Family Foundation, the Richard Gilder Graduate School at the American Museum of Natural History, and a Research Initiatives in Science and Engineering (RISE) award to PA from Columbia University. CSM was funded by a Macquarie University Research Excellence scholarship, a Fulbright Future Fellowship, and a Holsworth Wildlife Research Endowment from Equity Trustees Charitable Foundation and the Ecological Society of Australia. MK was supported by ERC POC Grant #101069216, HAPLOTAGGING awarded to YFC. YFC was supported by the Max Planck Society. Tissue loans were generously provided by the Australian National Wildlife Collection. We wish to thank Andrew Blunsden, Skye Davis, Emma Greig, Eva van der Heijden, Laura Hurley, Kyle Kostrzewa, Elise McCarthy, and Clément Rul for their assistance with fieldwork. Col Roberts and Adam Hunter suggested finching sites in Western Australia. Kelsie Lopez helped with DNA extractions, Amanda Carpenter helped with carotenoid analysis, Andrew Vaughn provided advice on selective sweep analyses. Trevor Price and Vicens Vila-Coury provided excellent feedback on the manuscript.

## Author Contributions

DMH and SCG designed research; DMH, CSM, MJP, NMJ, MK, and NH performed research; DMH analyzed genomic data; MJP and NMJ analyzed carotenoid data; DMH contributed tables and figures; YFC, GEH, PA, and SCG provided critical resources; and DMH wrote the paper with input from all authors.

## Competing Interest Statement

The authors declare no competing interests.

---

**Fig. S1.**
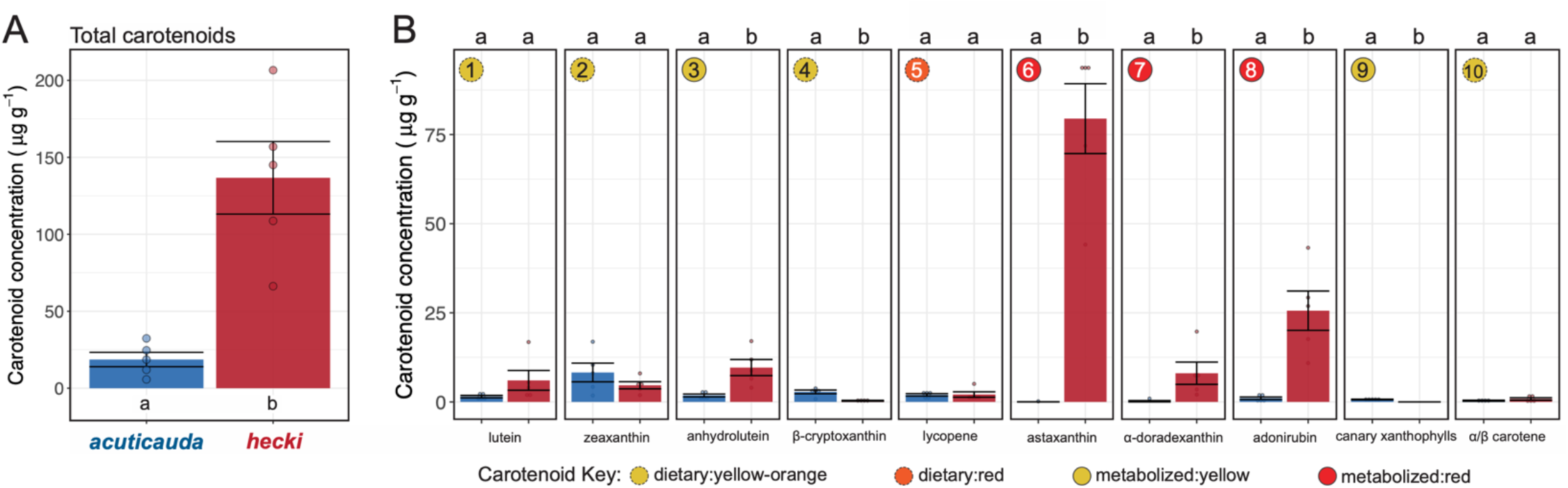
Bill integument carotenoid composition of each long-tailed finch subspecies. (A) Total carotenoid concentration in five individuals of subspecies *acuticauda* (blue) and *hecki* (red) as quantified via HPLC. (B) The concentration of ten distinct dietary and metabolized carotenoids observed in each subspecies. As in Fig. 2, each unique carotenoid has been numerically annotated from left to right and labeled in the top left of each panel as dietary yellow-orange (yellow dashed circle), dietary red (red dashed circle), metabolized yellow (solid yellow circle), or metabolized red (solid red circle). While the saponification method utilized in this study increases the recovery of ketocarotenoids and dietary carotenoids, it does result in the loss of canary xanthophylls a and b (metabolized yellow carotenoid #9) (Toomey et al. 2022b). As such, the inferred concentration of canary xanthophylls in the yellow bills of subspecies *acuticauda* is almost surely an underestimate. Boxplots represent subspecies means and whiskers represent subspecies mean and standard deviation, respectively. Notably, while canary xanthophylls were detected at low concentration in the yellow bills of subspecies *acuticauda*, no canary xanthophylls were detected in the red bills of subspecies *hecki*. The letters ‘a’ and ‘b’ above each plot indicate statistical significance between subspecies.

**Fig. S2.**
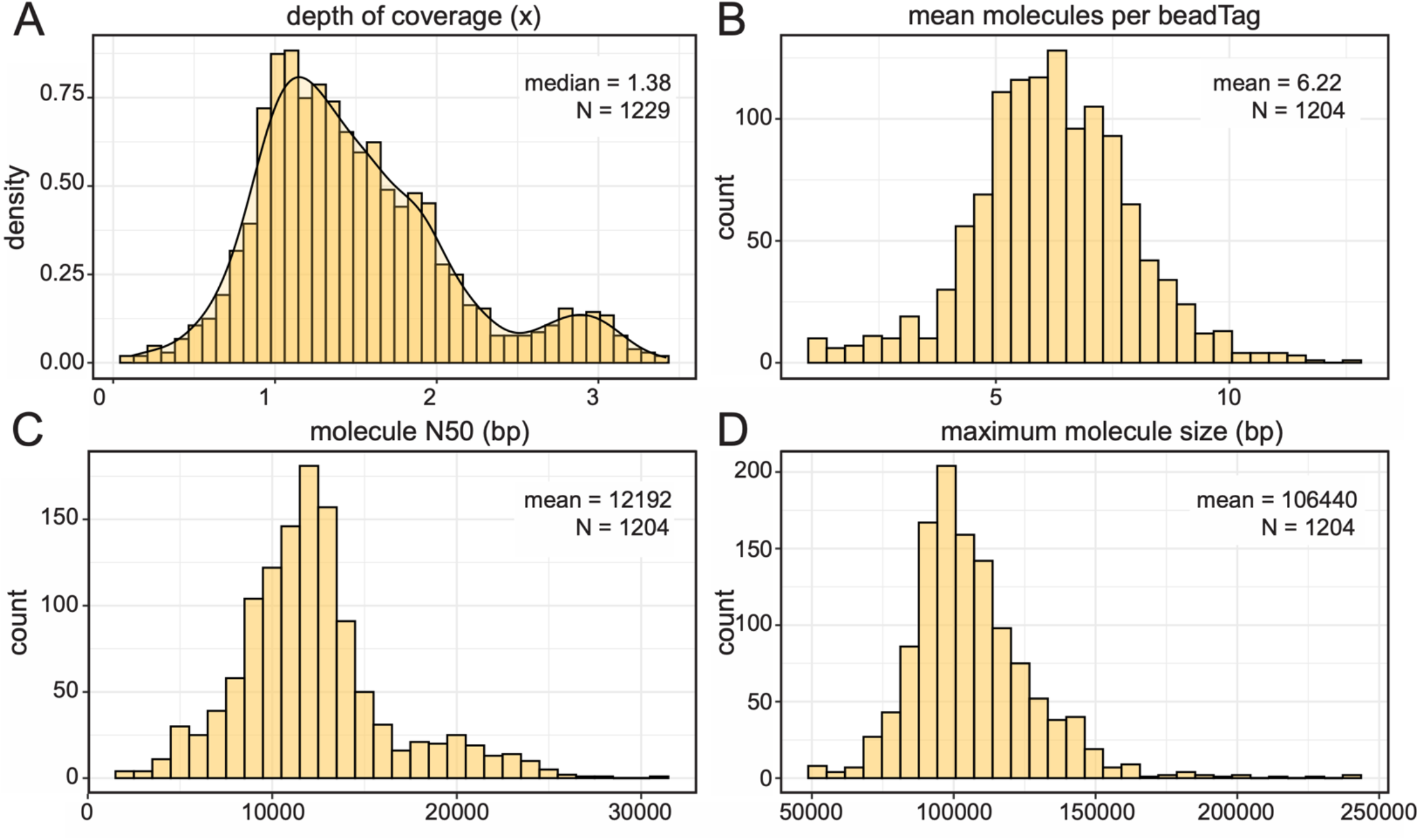
Sequencing effort and linked-read (LR) performance with haplotagging. (A) Depth of coverage as mapped to zebra finch reference bTaeGut1.4 (GCF_003957565.2) across 1133 long-tailed finch *Poephila acuticauda* and 96 black-throated finch *Poephila cincta* samples (N = 1229 samples total). (B) Distribution of per sample mean number of molecules per beadTag for 1204 samples with high molecular weight DNA (hmwDNA) available during LR library preparation. (C) Distribution of per sample molecule N50. (D) Distribution of per sample maximum molecule size.

**Fig. S3.**
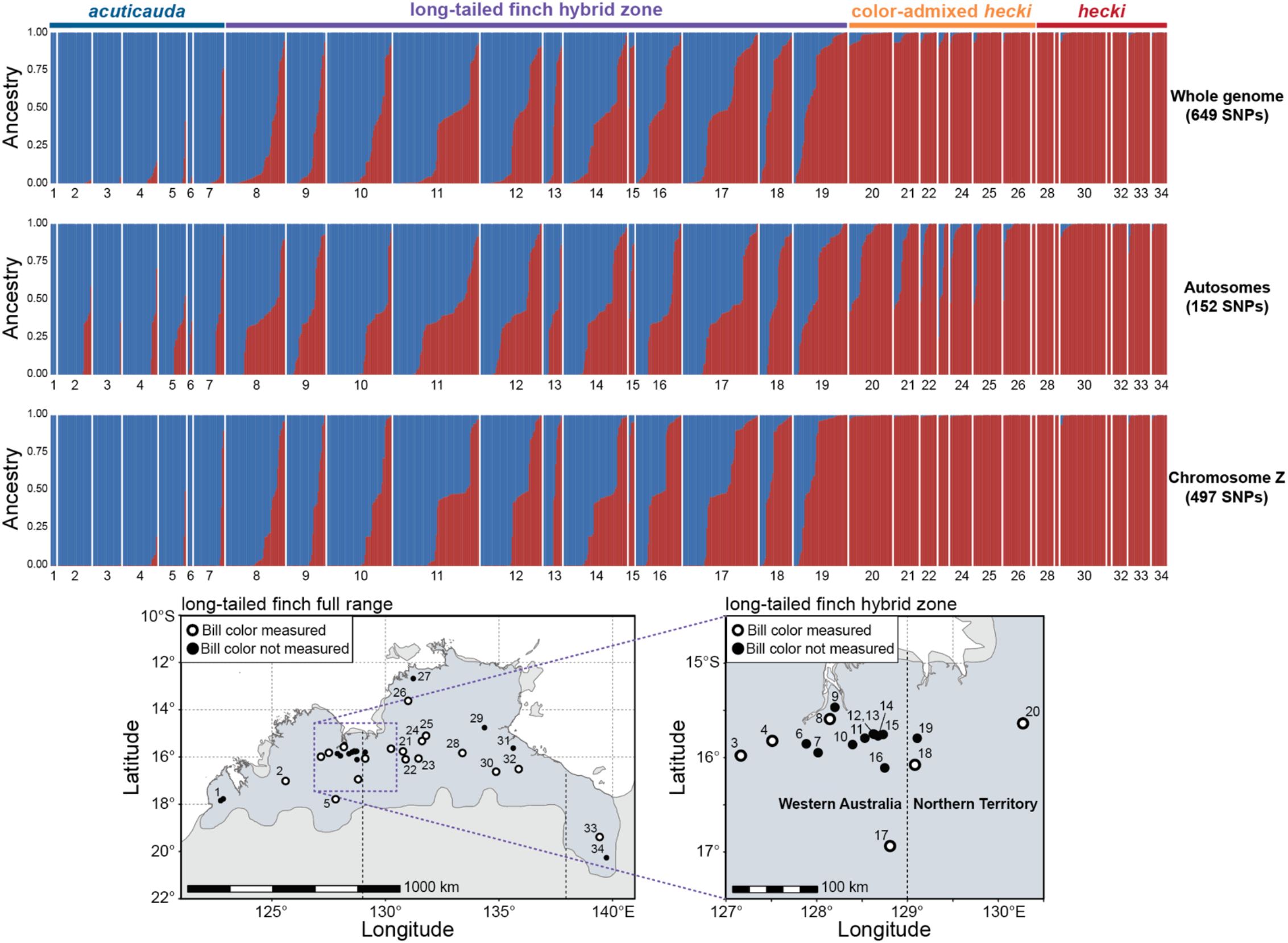
Genomic admixture between long-tailed finch subspecies *acuticauda* (blue) and *hecki* (red) is concentrated within a phenotypically cryptic hybrid zone between populations 8 and 19 located along the edge of the Kimberley Plateau on the western side of the Western Australia – Northern Territory border. Populations of color-admixed *hecki* (pops. 20-27) exhibit minimal *acuticauda* ancestry. Hybrid index scores were for 983 wild-sampled individuals were inferred with gghybrid using a set of LD-pruned ancestry informative markers genome-wide (N = 649 SNPs, top panel), on the autosomes (N = 152 SNPs, middle panel), and on the Z chromosome (N = 497 SNPs, bottom panel). The geographic location of each sampled population given in long-tailed finch range map and hybrid zone inset.

**Fig. S4.**
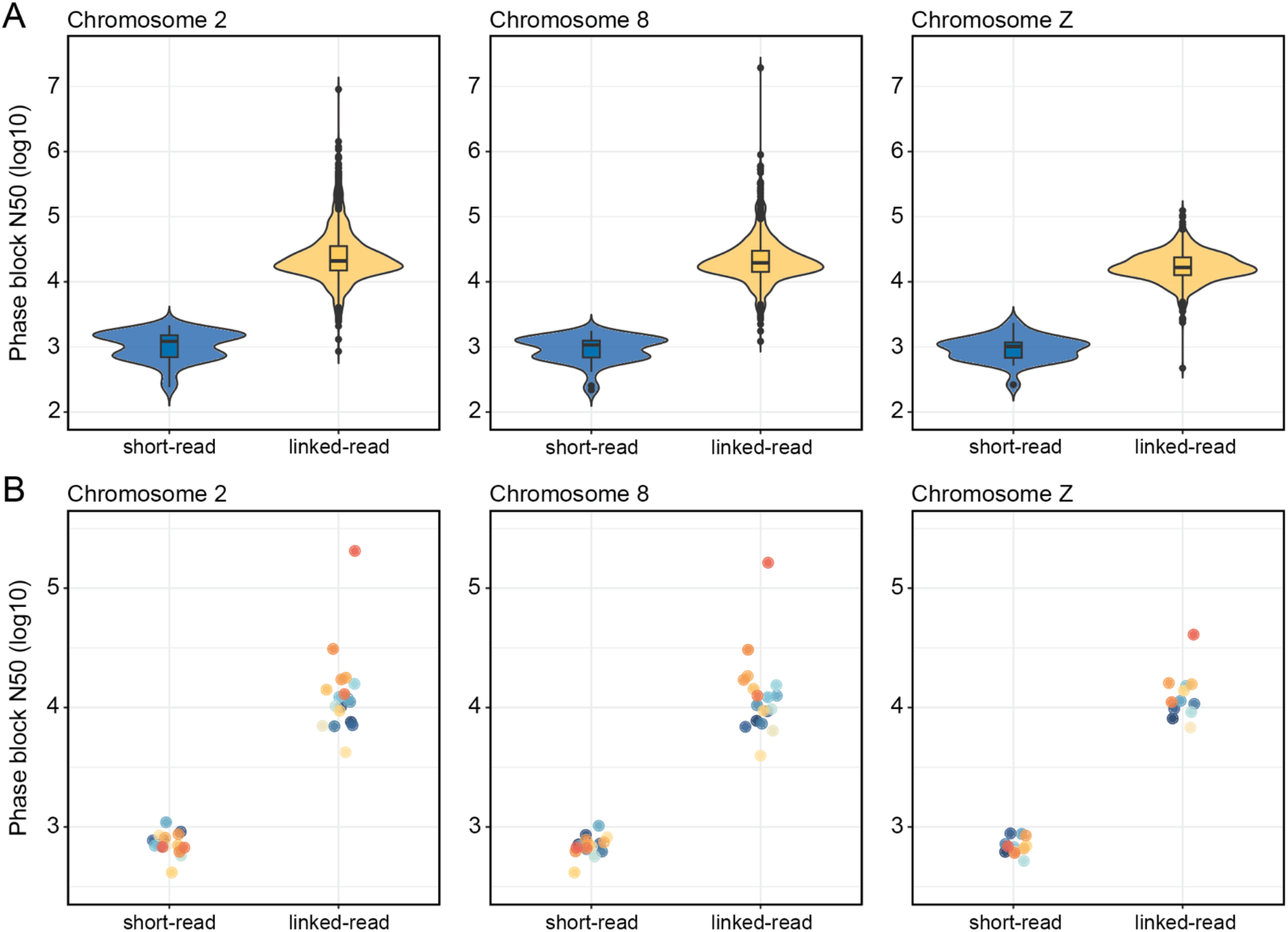
Comparison of phasing performance utilizing short-read (SR) and linked-read (LR) data. Phase block N50 calculated for 42 samples with SR WGS data and 1204 samples with LR data prepared using haplotagging chromosome 2, 8, and Z using HapCUT2 (Edge et al. 2017). (A) Distribution of phase block N50 by library type. Phasing performance substantially improved with LR data (median phase block N50, chr2 = 20.91 kbp, chr8 = 19.56 kbp, chrZ = 16.60 kbp) compared to phasing with SR data (median phase block N50, chr2 = 1.22 kbp, chr8 = 1.07 kbp, chrZ = 1.01 kbp). Only males used to evaluate phasing performance on chromosome Z (LR: N = 772, SR: N = 27). (B) Eighteen samples were available as technical replicates for evaluating phasing performance based on library type. Phase block N50s were on average two orders of magnitude larger when utilizing LR information (median phase block N50, chr2 = 11.57 kbp, chr8 = 11.32 kbp, chrZ = 11.25 kbp) compared to phasing with SR data (median phase block N50, chr2 = 0.70 kbp, chr8 = 0.69 kbp, chrZ = 0.68 kbp). Only 12 males were available as technical replicates for chromosome Z.

**Fig. S5.**
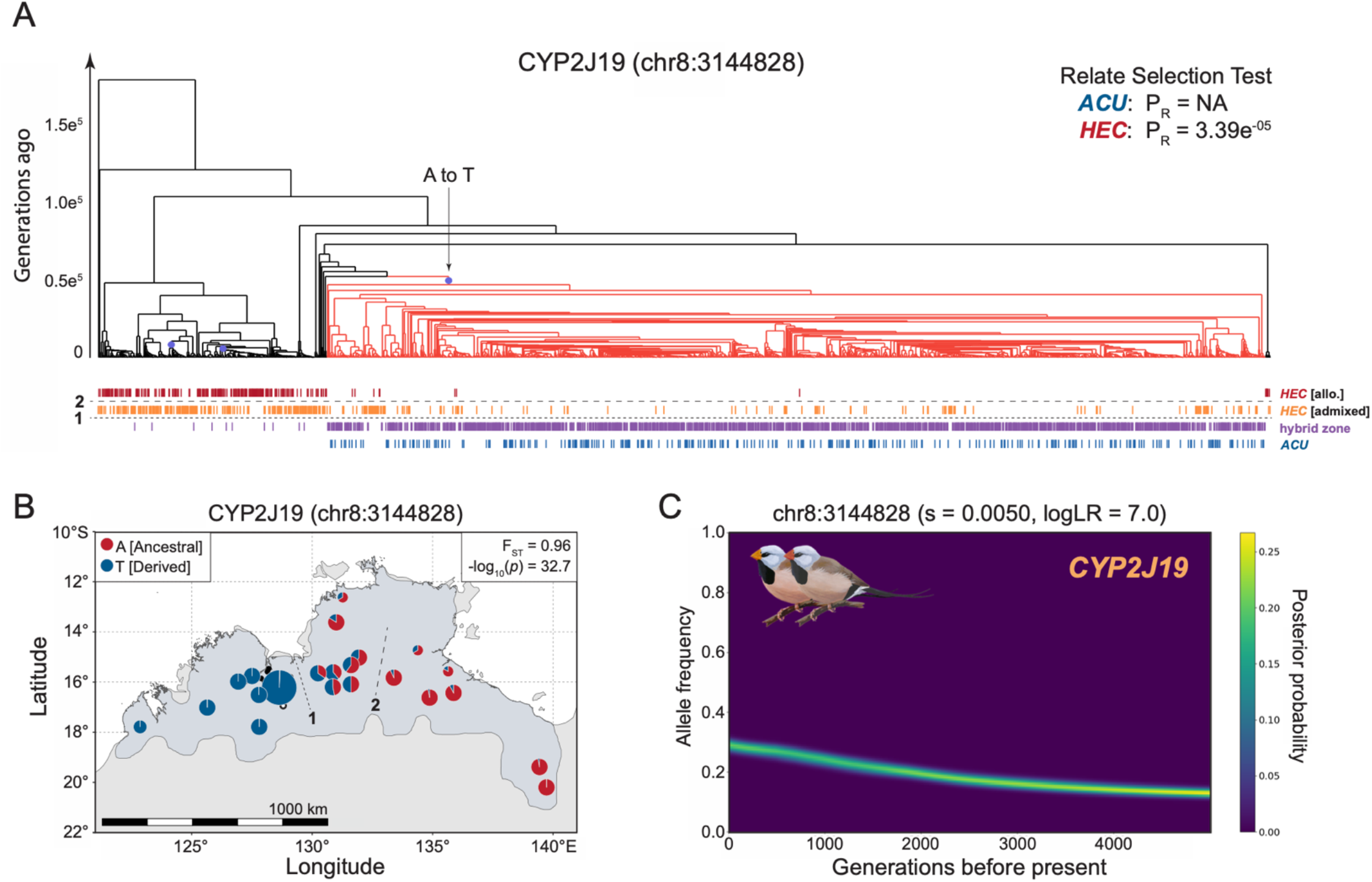
Evidence of selection on *CYP2J19* from ancestral recombination graph (ARG) inference is complicated by the age of the introgressing allele. (A) Relate marginal tree for the SNP most significantly associated with bill color variation: chr8:3144828 (2N = 1928 haplotypes). Branches colored in light red represent haplotypes carrying the derived allele at this site. Relate Selection Test evidence for selection over the lifetime of the mutation (P_R_) shown in top right for subspecies *acuticauda* (*ACU*: pops. 1-7, 2N = 282) and subspecies *hecki* (HEC: pops. 20-34, 2N = 522). Vertical hash marks beneath tips of marginal tree represent haplotypes from allopatric red-billed *hecki* (dark red, pops. 28-34), color-admixed *hecki* (orange, pops. 20-27), the hybrid zone (purple, pops. 8-19), and allopatric yellow-billed *acuticauda* (pops. 1-7). Horizontal dashed lines 1 and 2, as seen in panel (B), represent geographic breaks between the hybrid zone and color-admixed *hecki* (line 1) and between color-admixed and red-billed *hecki* (line 2). (B) Population allele frequencies for SNP chr8:3144828. Derived and ancestral alleles are color-coded blue and red, respectively. F_ST_ and GWAS significance given in top right inset of panel. (C) Historical allele trajectory of the *acuticauda*-derived allele at chr8:3144828 in subspecies *hecki*, which is associated with carotenoid ketolation enzyme *CYP2J19*. s, selection coefficient.

**Fig. S6.**
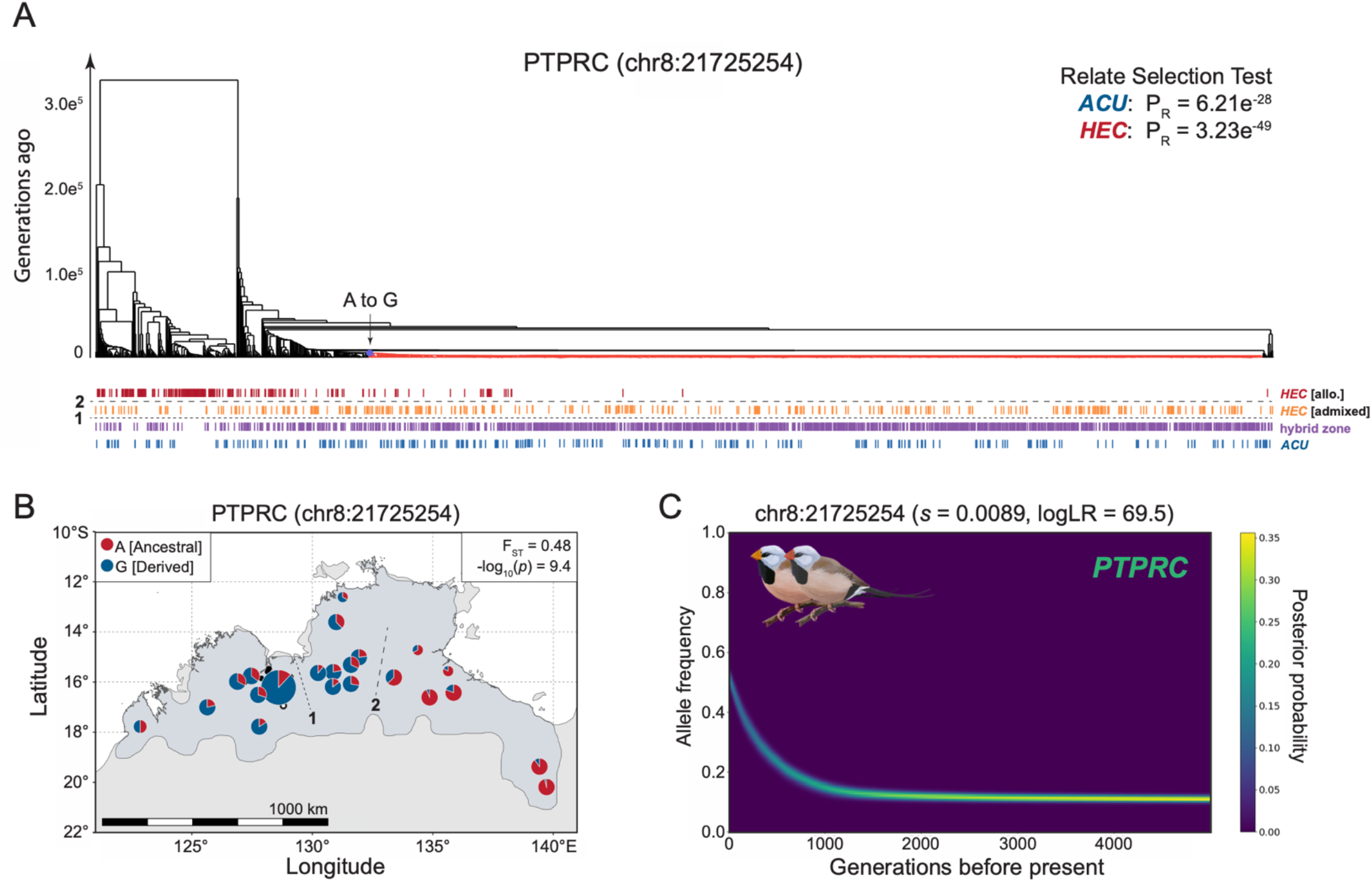
Evidence of selection on *PTPRC* from ancestral recombination graph (ARG) inference. (A) Relate marginal tree for bill color variation associated SNP chr8:21725254 (2N = 1928 haplotypes). Branches colored in light red represent haplotypes carrying the derived allele at this site. Relate Selection Test evidence for selection over the lifetime of the mutation (P_R_) shown in top right for subspecies *acuticauda* (*ACU*: pops. 1-7, 2N = 282) and subspecies *hecki* (HEC: pops. 20-34, 2N = 522). Vertical hash marks beneath tips of marginal tree represent haplotypes from allopatric red-billed *hecki* (dark red, pops. 28-34), color-admixed *hecki* (orange, pops. 20-27), the hybrid zone (purple, pops. 8-19), and allopatric yellow-billed *acuticauda* (pops. 1-7). Horizontal dashed lines 1 and 2, as seen in panel (B), represent geographic breaks between the hybrid zone and color-admixed *hecki* (line 1) and between color-admixed and red-billed *hecki* (line 2). (B) Population allele frequencies for SNP chr8:21725254. Derived and ancestral alleles are color-coded blue and red, respectively. F_ST_ and GWAS significance given in top right inset of panel. (C) Historical allele trajectory of the *acuticauda*-derived allele at chr8:21725254 in subspecies *hecki*, which is associated with the tyrosine phosphatase receptor *PTPRC*. s, selection coefficient.

**Fig. S7.**
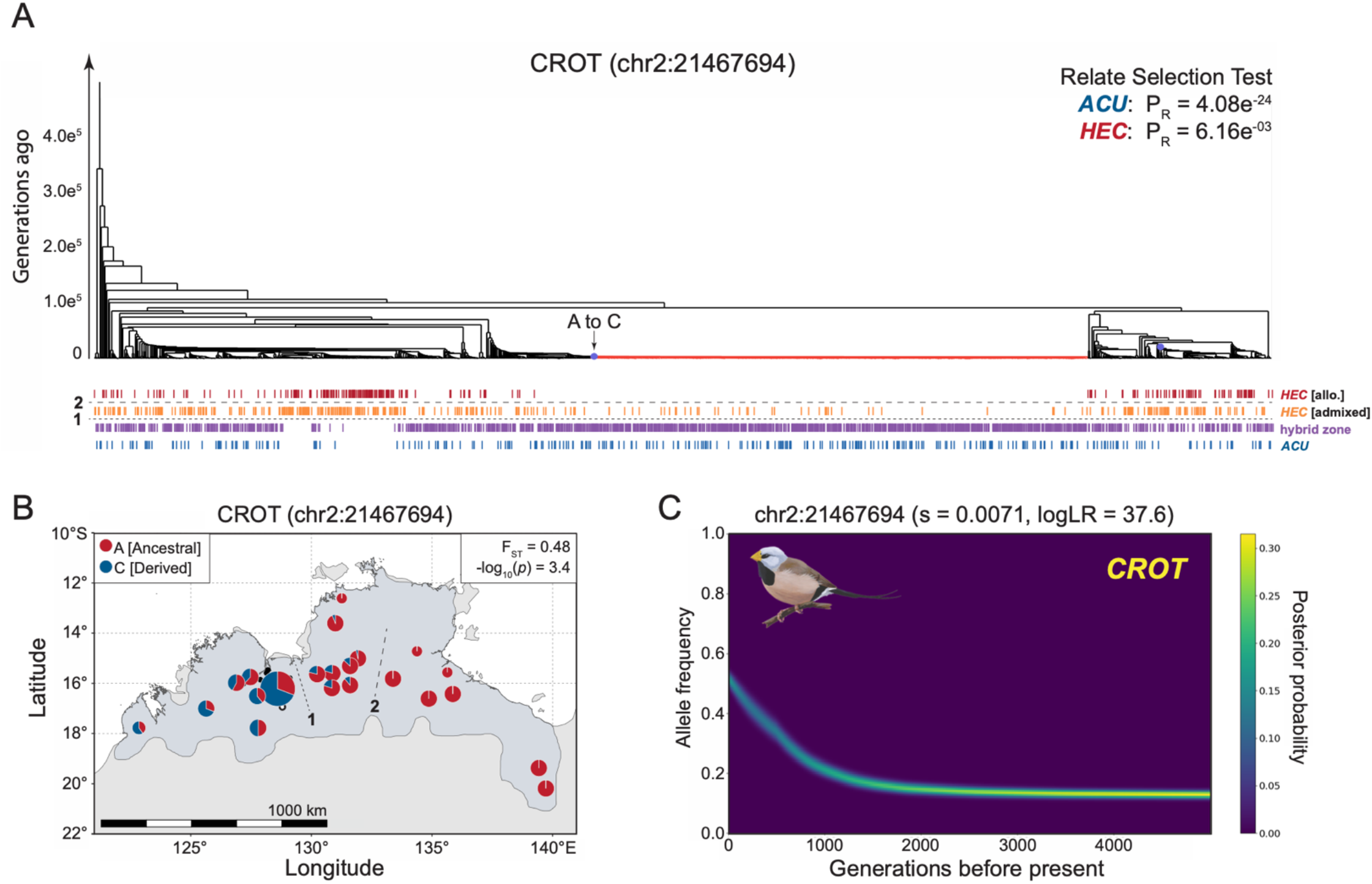
Evidence of selection on *CROT* from ancestral recombination graph (ARG) inference. (A) Relate marginal tree for bill color variation associated SNP chr2:21467694 (2N = 1928 haplotypes). Branches colored in light red represent haplotypes carrying the derived allele at this site. Relate Selection Test evidence for selection over the lifetime of the mutation (P_R_) shown in top right for subspecies *acuticauda* (*ACU*: pops. 1-7, 2N = 282) and subspecies *hecki* (HEC: pops. 20-34, 2N = 522). Vertical hash marks beneath tips of marginal tree represent haplotypes from allopatric red-billed *hecki* (dark red, pops. 28-34), color-admixed *hecki* (orange, pops. 20-27), the hybrid zone (purple, pops. 8-19), and allopatric yellow-billed *acuticauda* (pops. 1-7). Horizontal dashed lines 1 and 2, as seen in panel (B), represent geographic breaks between the hybrid zone and color-admixed *hecki* (line 1) and between color-admixed and red-billed *hecki* (line 2). (B) Population allele frequencies for SNP chr2:21467694. Derived and ancestral alleles are color-coded blue and red, respectively. F_ST_ and GWAS significance given in top right inset of panel. (C) Historical allele trajectory of the acuticauda-derived allele at chr2:21467694 in subspecies acuticauda, which is associated with mitochondrial fatty acid β-oxidation enzyme *CROT*. s, selection coefficient.

**Fig S8.**
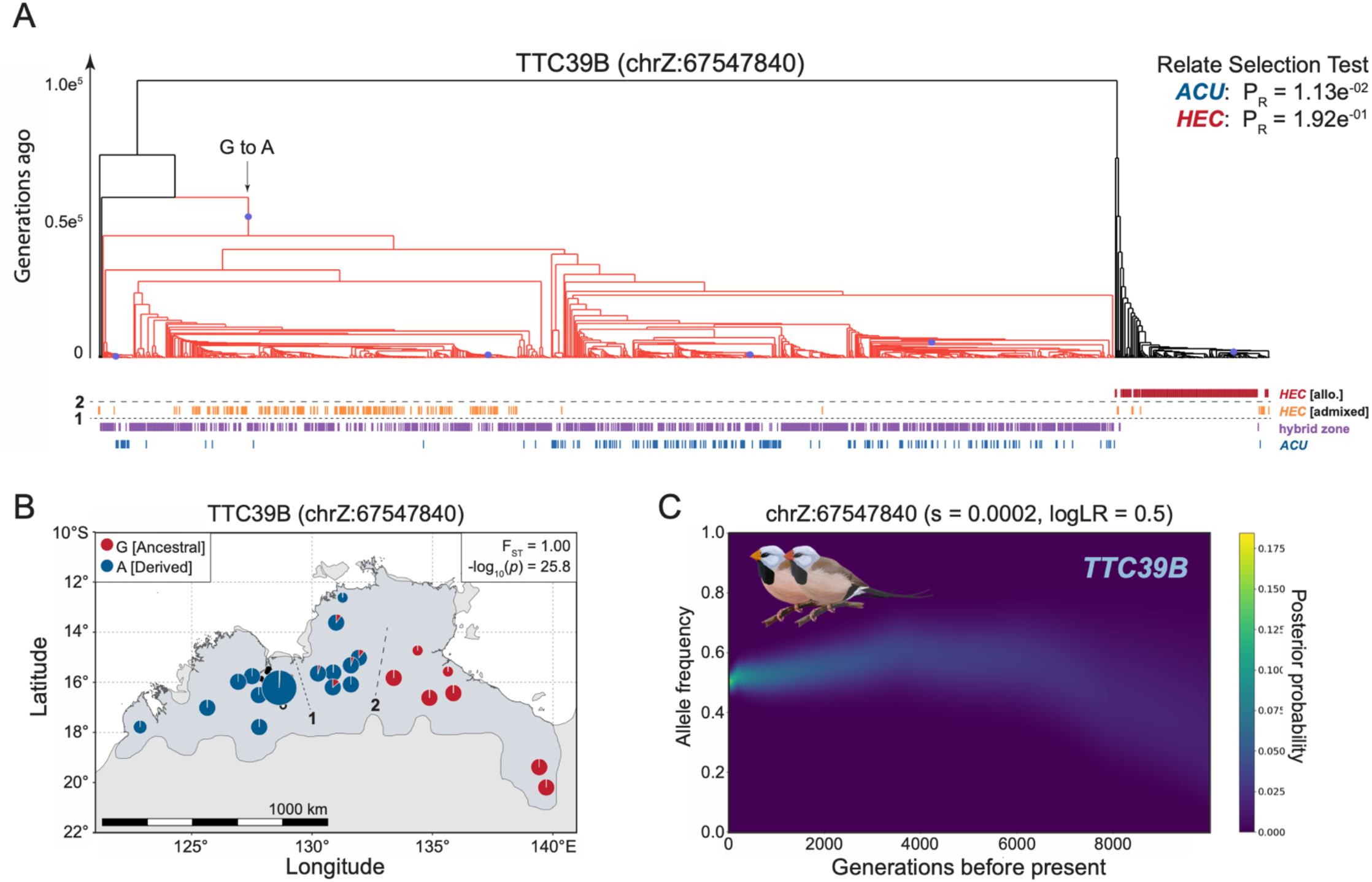
Evidence of selection on *TTC39B* from ancestral recombination graph (ARG) inference is complicated by the age of the introgressing allele. (A) Relate marginal tree for the second most significant SNP associated with bill color variation: chrZ:67547840 (2N = 1206 haplotypes). Branches colored in light red represent haplotypes carrying the derived allele at this site. Relate Selection Test evidence for selection over the lifetime of the mutation (P_R_) shown in top right for subspecies *acuticauda* (*ACU*: pops. 1-7, 2N = 178) and subspecies *hecki* (HEC: pops. 20-34, 2N = 312). Vertical hash marks beneath tips of marginal tree represent haplotypes found in males (i.e., with two copies of the Z chromosome) from allopatric red-billed *hecki* (dark red, pops. 28-34), color-admixed *hecki* (orange, pops. 20-27), the hybrid zone (purple, pops. 8-19), and allopatric yellow-billed *acuticauda* (pops. 1-7). Horizontal dashed lines 1 and 2, as seen in panel (B), represent geographic breaks between the hybrid zone and color-admixed *hecki* (line 1) and between color-admixed and red-billed *hecki* (line 2). (B) Population allele frequencies for SNP chrZ:67547840. Derived and ancestral alleles are color-coded blue and red, respectively. F_ST_ and GWAS significance given in top right inset of panel. (C) Historical allele trajectory of the derived allele at chrZ: 67547840 in subspecies *hecki*, which is located within an intron of carotenoid ketolation enhancer gene *TTC39B*. s, selection coefficient.

**Fig. S9.**
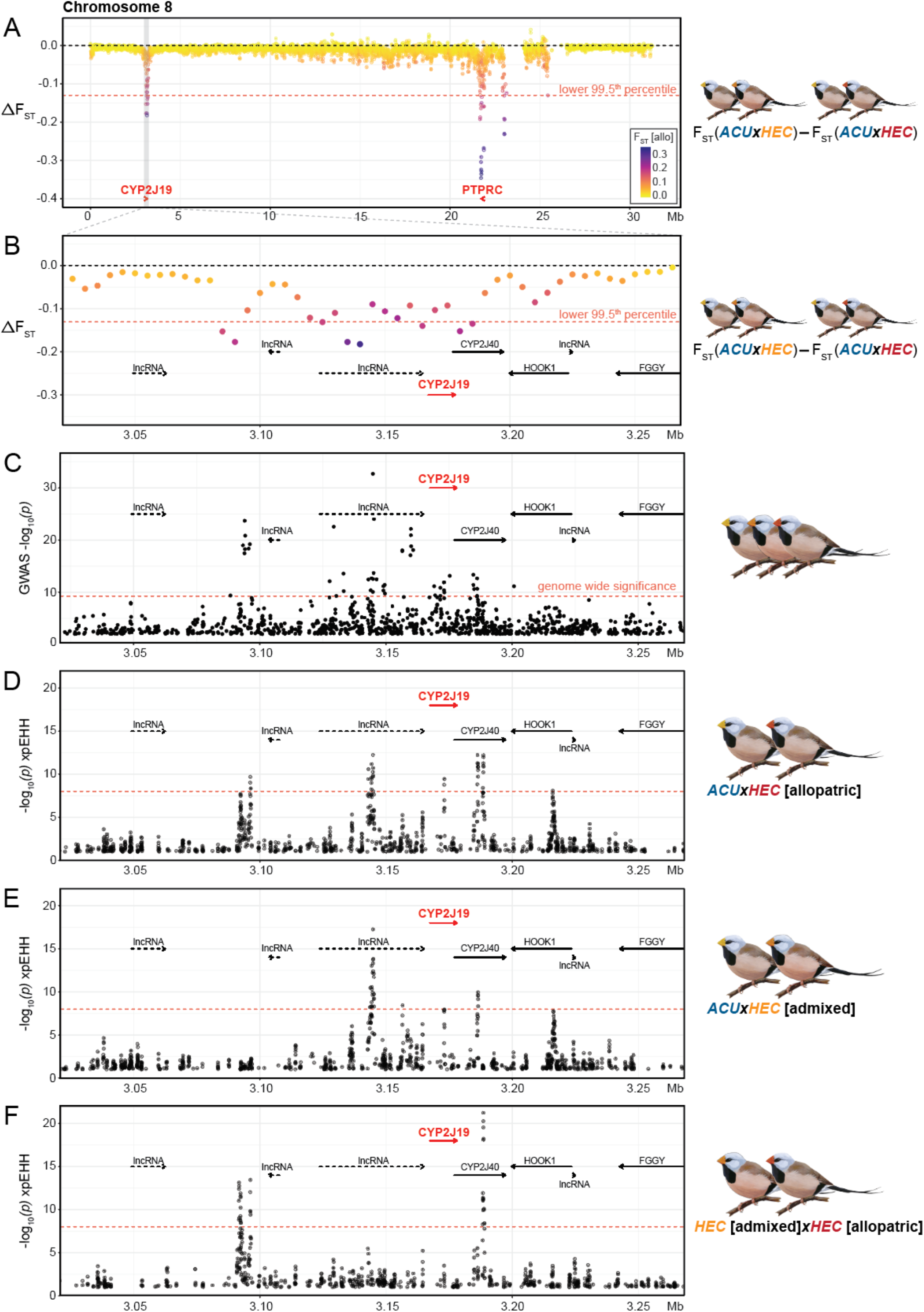
Evidence of introgression and selection on chromosome 8 associated with the carotenoid ketolation enzyme *CYP2J19*. (A) Chromosome wide ΔF_ST_ in 10 kb windows with 5 kb step size. Points are color-coded by F_ST_ between allopatric populations of each subspecies. ΔF_ST_ outliers below the 99.5^th^ percentile – denoted as a horizontal dashed red line – represent regions that have introgressed from *acuticauda* into *hecki*. The location of *CYP2J19* and *PTPRC* are shown as red arrows. (B) Zoom-in of the introgression outlier window encompassing *CYP2J19*. (C) GWAS results for SNPs within this window. (D to F) Cross-population extended haplotype homozygosity (xpEHH) statistical significance between allopatric *acuticauda* and allopatric *hecki* (D), between allopatric *acuticauda* and color-admixed *hecki* (E), and between color-admixed *hecki* and allopatric *hecki* (F).

**Fig. S10.**
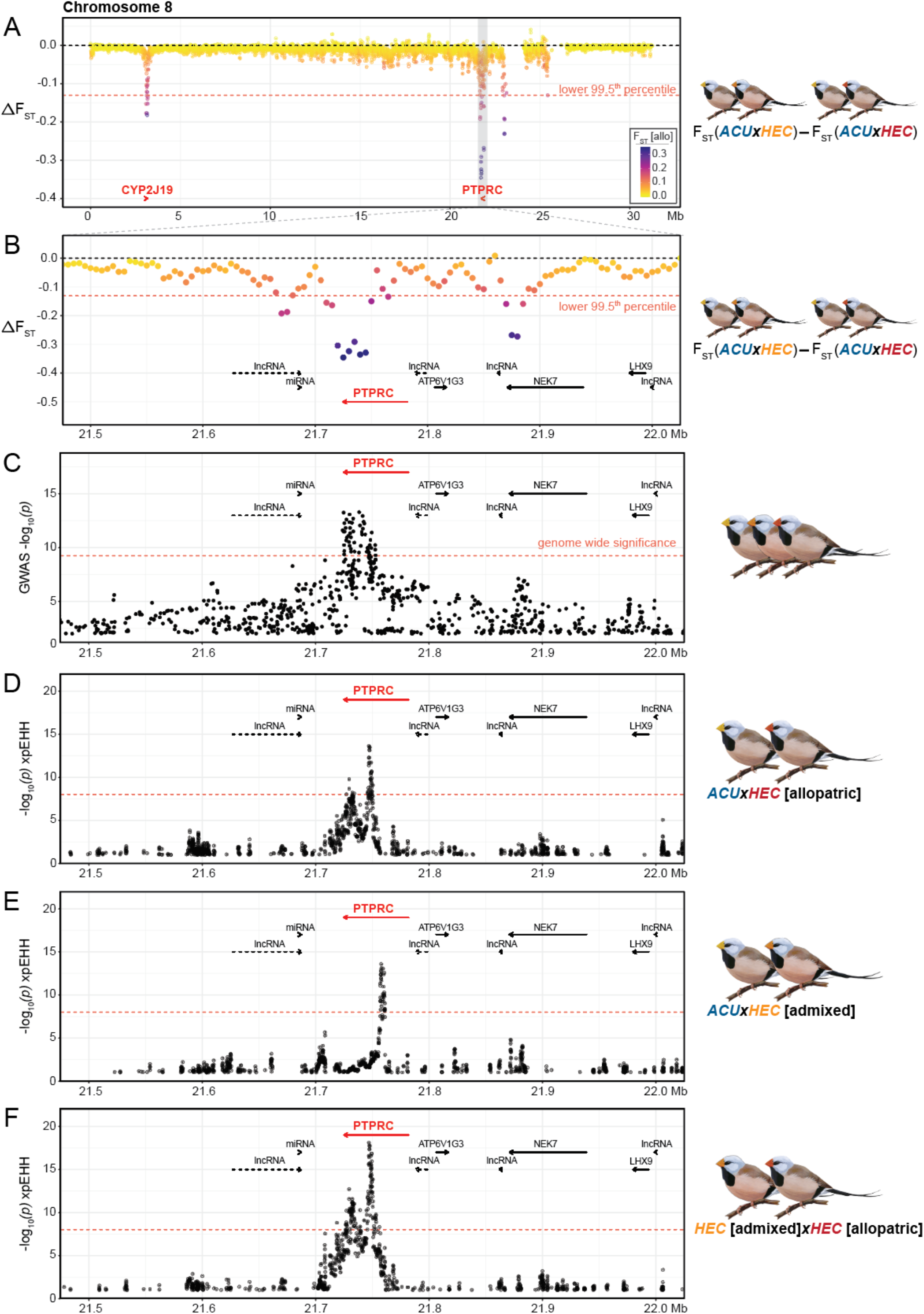
Evidence of introgression and selection on chromosome 8 associated with the tyrosine phosphatase receptor *PTPRC*. (A) Chromosome wide ΔF_ST_ in 10 kb windows with 5 kb step size. Points are color-coded by F_ST_ between allopatric populations of each subspecies. ΔF_ST_ outliers below the 99.5^th^ percentile – denoted as a horizontal dashed red line – represent regions that have introgressed from *acuticauda* into *hecki*. The location of *CYP2J19* and *PTPRC* are shown as red arrows. (B) Zoom-in of the introgression outlier window encompassing *PTPRC*. (C) GWAS results for SNPs within this window. (D to F) Cross-population extended haplotype homozygosity (xpEHH) statistical significance between allopatric *acuticauda* and allopatric *hecki* (D), between allopatric *acuticauda* and color-admixed *hecki* (E), and between color-admixed *hecki* and allopatric *hecki* (F).

**Fig. S11.**
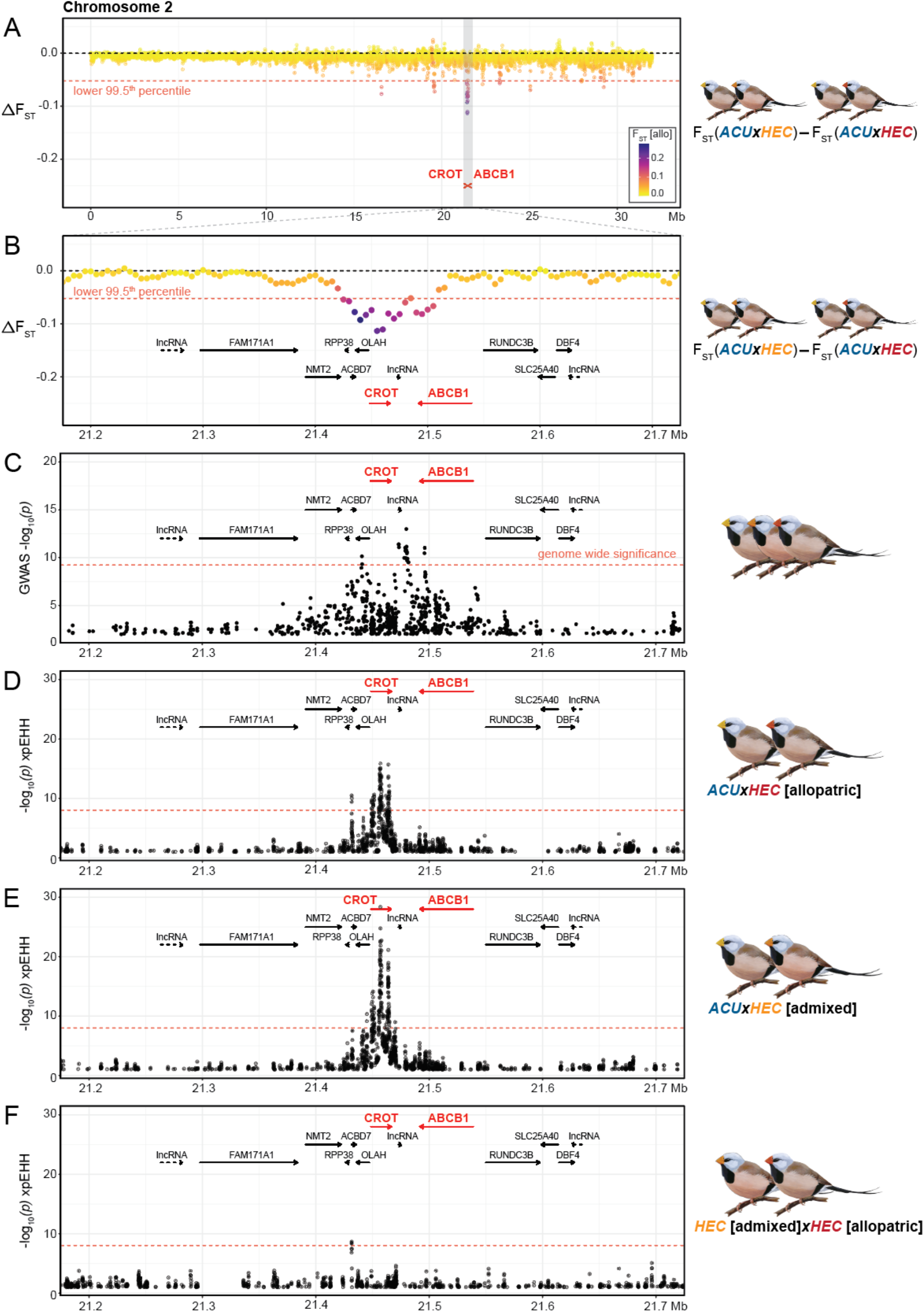
Evidence of introgression and selection on chromosome 2 associated with mitochondrial fatty acid *β*-oxidation enzyme *CROT*. (A) Chromosome wide ΔF_ST_ in 10 kb windows with 5 kb step size (chromosome 2 is shown truncated from 0 to 35 Mb for clarity). Points are color-coded by F_ST_ between allopatric populations of each subspecies. ΔF_ST_ outliers below the 99.5^th^ percentile – denoted as a horizontal dashed red line – represent regions that have introgressed from *acuticauda* into *hecki*. The location of *CROT* and *ABCB1* are shown as red arrows. (B) Zoom-in of the introgression outlier window encompassing *CROT* and *ABCB1*. (C) GWAS results for SNPs within this window. (D to F) Cross-population extended haplotype homozygosity (xpEHH) statistical significance between allopatric *acuticauda* and allopatric *hecki* (D), between allopatric *acuticauda* and color-admixed *hecki* (E), and between color-admixed *hecki* and allopatric *hecki* (F).

**Fig. S12.**
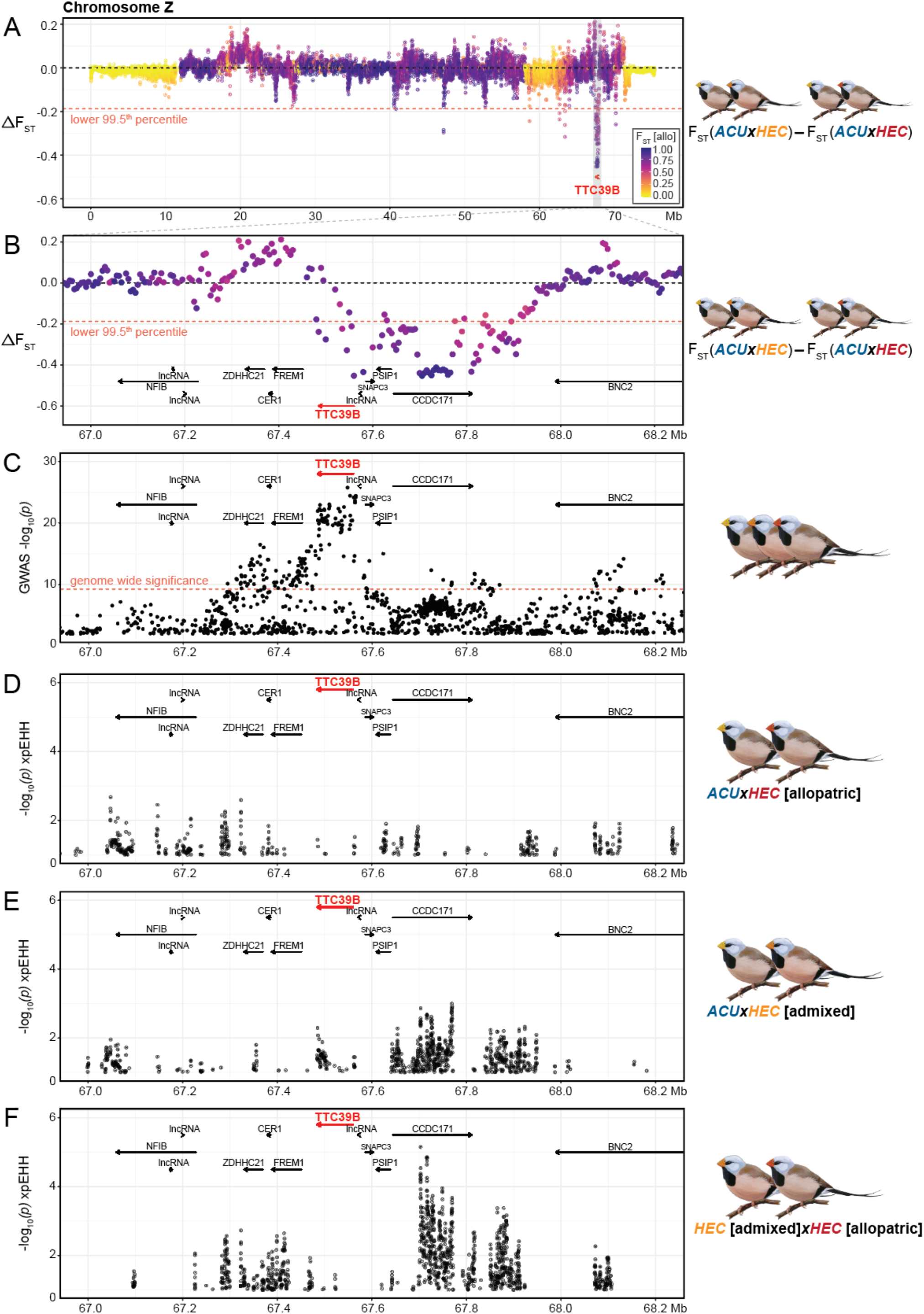
Evidence of introgression and selection on chromosome Z associated with carotenoid ketolation enhancer gene *TTC39B*. (A) Chromosome wide ΔF_ST_ in 10 kb windows with 5 kb step size. Points are color-coded by F_ST_ between allopatric populations of each subspecies. ΔF_ST_ outliers below the 99.5^th^ percentile – denoted as a horizontal dashed red line – represent regions that have introgressed from *acuticauda* into *hecki*. The location of *TTC39B* is shown as a red arrow. (B) Zoom-in of the introgression outlier window encompassing *TTC39B*. (C) GWAS results for SNPs within this window. (D to F) Cross-population extended haplotype homozygosity (xpEHH) statistical significance between allopatric *acuticauda* and allopatric *hecki* (D), between allopatric *acuticauda* and color-admixed *hecki* (E), and between color-admixed *hecki* and allopatric *hecki* (F).

**Fig S13.**
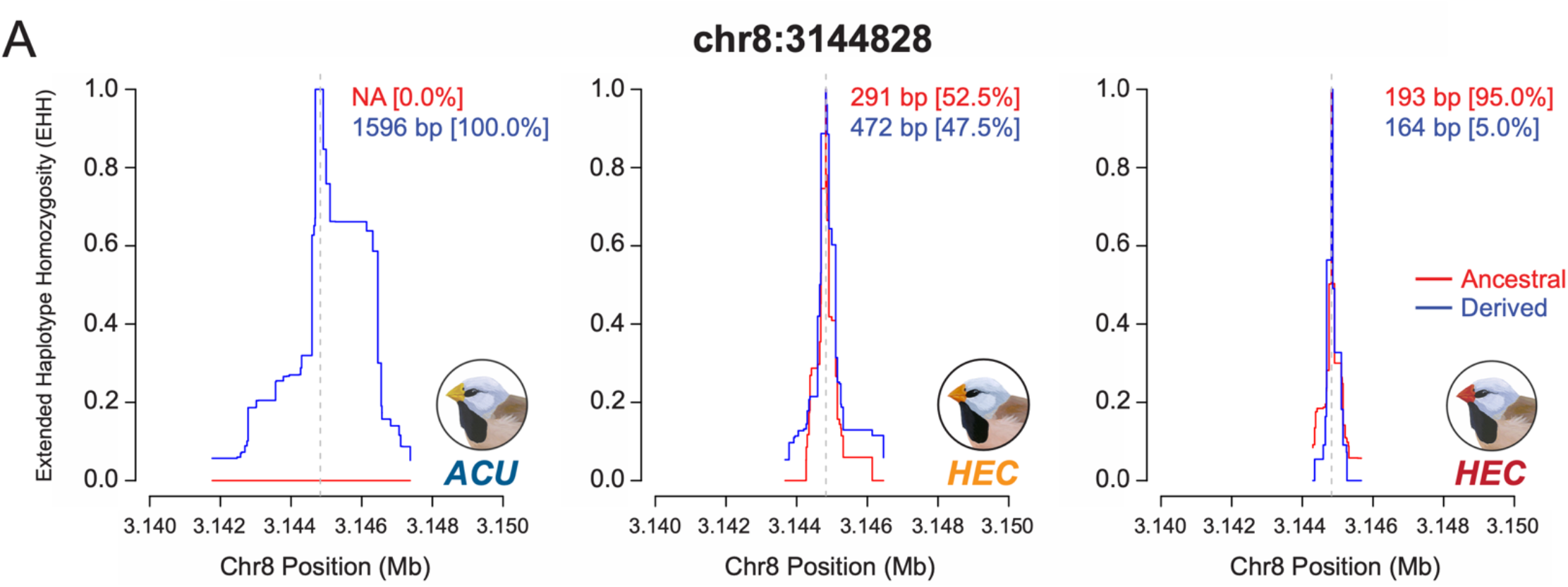
Extended haplotype homozygosity (EHH) summaries for bill hue associated SNP chr8:3144828, located ~20 kbp upstream of *CYP2J19*, which has xpEHH support of a selective sweep in *acuticauda* relative to both color-admixed (-log10(P) = 17.3) and red-billed *hecki* (-log10(P) = 9.9). From left, EHH was calculated using phased haplotype data from *acuticauda* (pops. 1 – 7, 2N = 276), color-admixed *hecki* (pops. 20 – 27, 2N = 320), and red-billed *hecki* (pops. 28 – 34, 2N = 202).

**Fig S14.**
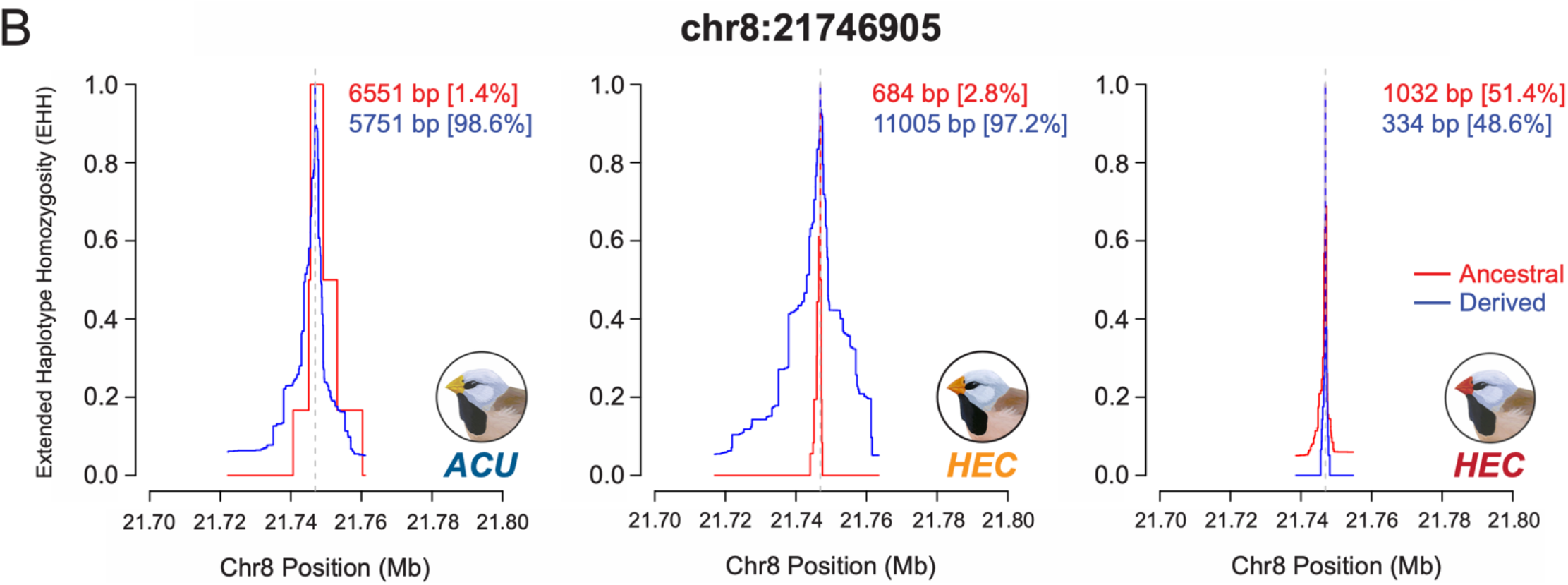
Extended haplotype homozygosity (EHH) summaries for bill hue associated SNP chr8:21746905, located within an intron of *PTPRC*, which has xpEHH support of a selective sweep in color-admixed *hecki* relative to red-billed *hecki* (-log10(P) = 18.1). From left, EHH was calculated using phased haplotype data from *acuticauda* (pops. 1 – 7, 2N = 276), color-admixed *hecki* (pops. 20 – 27, 2N = 320), and red-billed *hecki* (pops. 28 – 34, 2N = 202).

**Fig. S15.**
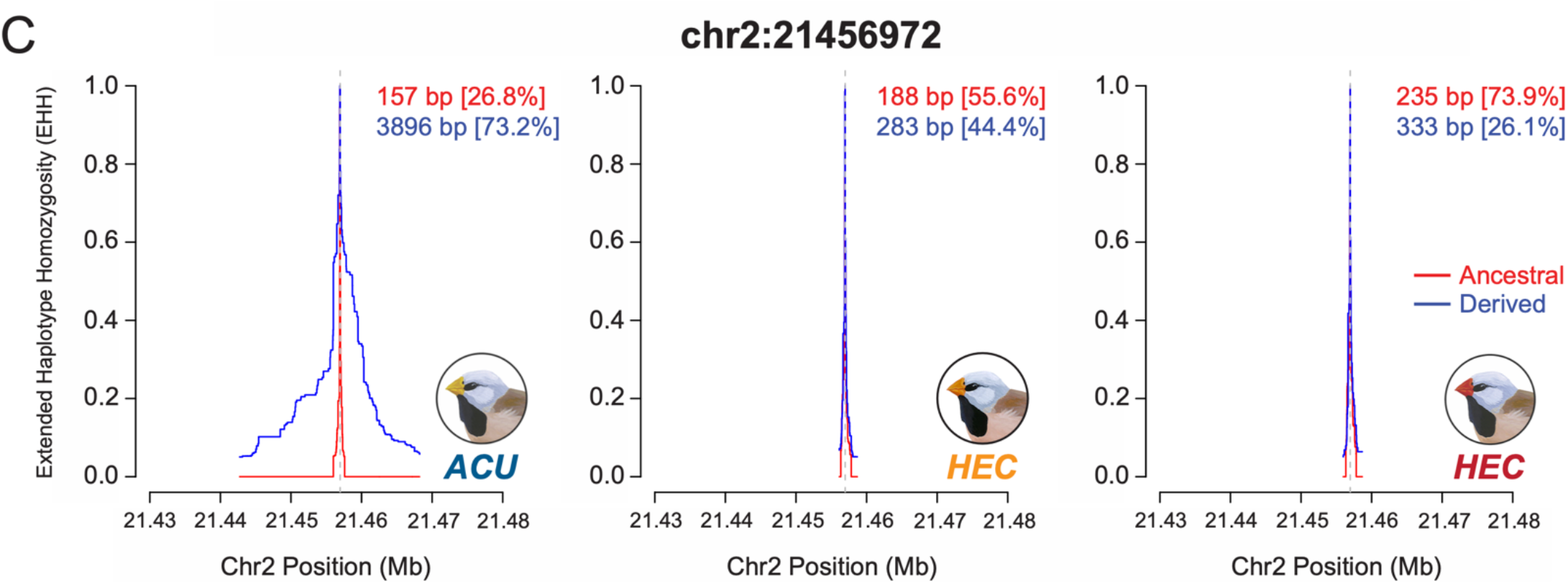
Extended haplotype homozygosity (EHH) summaries for bill hue associated SNP chr2:21456972, located within an intron of *CROT*, which has xpEHH support of a selective sweep in *acuticauda* relative to both color-admixed (-log10(P) = 28.4) and red-billed *hecki* (-log10(P) = 15.9). From left, EHH was calculated using phased haplotype data from *acuticauda* (pops. 1 – 7, 2N = 276), color-admixed *hecki* (pops. 20 – 27, 2N = 320), and red-billed *hecki* (pops. 28 – 34, 2N = 202).

**Fig. S16.**
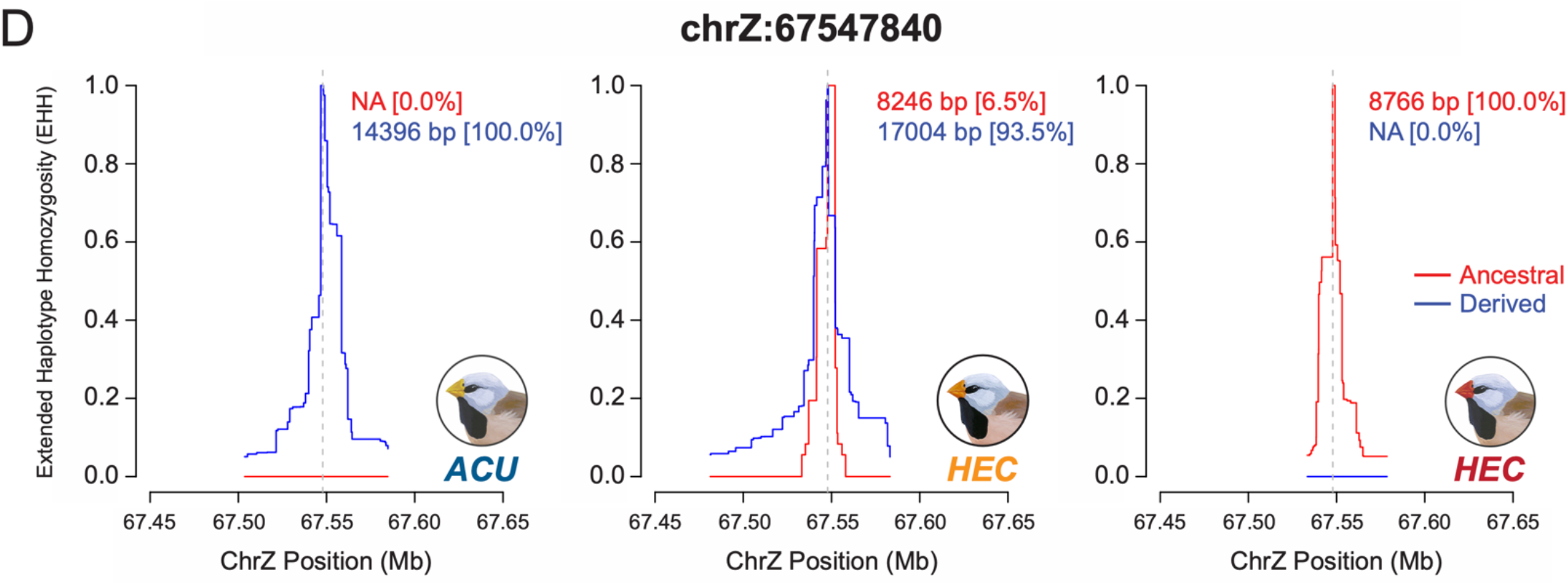
Extended haplotype homozygosity (EHH) summaries for bill hue associated SNP chrZ:67547840, located within an intron of *TTC39B*. This locus demonstrates the difficulty of calculating xpEHH between focal populations due to their difference in allele frequencies. In populations of color-admixed *hecki*, center panel, haplotype homozygosity is >2× greater for carriers of the *acuticauda*-derived variant at this locus, consistent with a selective sweep following introgression across the long-tailed finch hybrid zone. From left, EHH was calculated using phased haplotype data from males in *acuticauda* (pops. 1 – 7, 2N = 160), color-admixed *hecki* (pops. 20 – 27, 2N = 168), and red-billed *hecki* (pops. 28 – 34, 2N = 128).

## Notes

### Competing Interest Statement

The authors have declared no competing interest.

